# A direct role for SNX9 in the biogenesis of filopodia

**DOI:** 10.1101/710285

**Authors:** IK Jarsch, JR Gadsby, A Nuccitelli, J Mason, H Shimo, L Pilloux, B Marzook, CM Mulvey, U Dobramysl, KS Lilley, RD Hayward, TJ Vaughan, CL Dobson, JL Gallop

## Abstract

Filopodia are finger-like actin-rich protrusions that extend from the cell surface and are important for cell-cell communication and pathogen internalization. The small size and transient nature of filopodia combined with shared usage of actin regulators within cells confounds attempts to identify filopodial proteins. Here, we used phage display phenotypic screening to isolate antibodies that alter the actin morphology of filopodia-like structures *in vitro*. We found that all of the antibodies that cause shorter FLS interact with SNX9, an actin regulator that binds phosphoinositides during endocytosis and in invadopodia. In cells, we discover SNX9 at specialised filopodia in *Xenopus* development and that SNX9 is an endogenous component of filopodia that are hijacked by *Chlamydia* entry. We show the use of antibody technology to identify proteins used in filopodia-like structures, and a role for SNX9 in filopodia.

## Introduction

Assemblies of actin filaments (F-actin) are major dynamic superstructures required for cell motility, inter-cell communication, and force generation. Filopodia contain parallel, tightly bundled arrays of F-actin and protrude from the cell to traverse substantial distances to detect substrate stiffness, the extracellular matrix and initiate cellular responses to growth factors. Filopodia are hijacked by bacterial and viral pathogens during cell entry and can also facilitate cell-to-cell transmission (Chang et al., 2016)They are a precursor structure in the formation of synapses and dendritic spines (Gallo, 2013), while specialised filopodia called cytonemes deliver morphogens to map the arrangements of tissues (Mattes and Scholpp, 2018).

What we know so far about how filopodia form remains poorly understood: while filopodia appear to arise from Arp2/3-dependent actin polymerisation (Yang and Svitkina, 2011), additional nucleation pathways, such as from formins, have also been implicated (Faix et al., 2009). Ena/VASP proteins are important in filopodia formation in cortical neurons and terminal arborisation in retinal ganglion cells, while in osteosarcoma cells although reducing levels of Ena/VASP proteins inhibits filopodia, Ena/VASP localize to focal adhesions in the cell body rather than at filopodia tips (Young et al., 2018). The actin bundling protein fascin is a major component of filopodia, however the spacing of F-actin is more consistent with cross-linking by the alternative actin bundling protein fimbrin (Jasnin et al., 2013). Myosin-X clusters at filopodial tips where it recruits a range of filopodial and adhesion proteins and new actin monomers are incorporated (He et al., 2017). While actin disassembly conventionally would occur by ADF/cofilin-mediated treadmilling at the pointed end of the filament, cofilin can synergise with fascin to induce actin disassembly at filopodial tips (Breitsprecher et al., 2011). Despite the fundamental importance of filopodia in neuronal networks, host-pathogen interactions and wider developmental processes, to date there is no unifying model to explain how the actin cytoskeleton and membrane machinery are coupled to initiate, elongate and sustain the dynamic behaviour of filopodia. Analysis of filopodial biogenesis is complicated since the actin regulatory proteins involved have overlapping roles in cell adhesion and endocytosis.

Many of these limitations associated with studying actin dynamics in cells can be circumvented by exploiting cell-free systems using cytoplasmic extracts. Cell-free approaches allow the functional reconstitution of defined actin-dependent processes and the biochemical analysis of individual components in response to defined signals. We use a cell-free system to reconstitute filopodia-like structures (FLS) that employs high-speed supernatent *Xenopus* egg extracts and supported lipid bilayers of defined and controllable composition as a cellular bilayer mimetic. Dynamic FLS extend from a surface of supported lipid bilayers augmented with phosphoinositol (4,5) bisphosphate (Lee et al., 2010). Our previous work demonstrated that the initial stage of FLS growth is sensitive to inhibition of Arp2/3-dependent actin nucleation whereas steady-state growth dynamics involve an interchanging complement of actin elongation and bundling proteins including the formin Drf3 (mDia2), Ena, VASP and fascin (Dobramysl et al., 2019; Lee et al., 2010). There are striking similarities between FLS and filopodia: heterogeneous “tip complexes” of actin regulators assemble on the supported lipid bilayers, and both FLS and filopodia display stochastic growth and retraction with similar quantitative dynamics (Dobramysl et al., 2019). A mechanistic understanding of filopodia is therefore attainable using a combination of cell-free assays, reconstitution and cellular models.

For the *de novo* identification of the proteins involved in making FLS, here we have combined the FLS system with phage display phenotypic screening. In this approach, a large library of 1×10^11^ human single chain Fv fragments (scFv) displayed on the surface of bacteriophage was used to select antibodies engaged in specific interactions with different FLS. Since the gene encoding the specific scFv is contained within the bacteriophage, the nucleotide sequence of the variable region can be determined and the antibody expressed and purified. The antibodies are then used in a functional assay to select those that generate phenotypes of particular interest. Antigens can then be identified by use of protein microarrays or immunoprecipitation followed by mass spectrometry. While antibodies have much recognized success in perturbing function extracellularly as therapeutics and research tools, the FLS system enables access to intracellular antigens. Therefore exploiting the FLS system provides a powerful new way of using antibody technology to identify novel factors and molecular mechanisms of actin regulation.

Our screen has identified a range of antibodies that generate different phenotypes in the FLS system. We have identified one of the antigens as SNX9, a known actin regulatory protein involved in invadopodia and endocytosis that interacts with phosphoinositide lipids and dynamin. Additionally, based on this finding, we have discovered roles for dynamin and phosphoinositide lipid metabolism in the growth of FLS. Furthermore, we observe that SNX9 directly localises to specialized filopodia in dorsal marginal zone explants from *Xenopus* embryos, where filopodia are used in Wnt signalling (Mattes et al., 2018). In cultured mammalian cells, we find SNX9 localizes to the tip and shaft almost all filopodia, providing evidence for a direct role of SNX9 in the filopodial entry mechanism used by *Chlamydia trachomatis* (Ford et al., 2018).

## Result

### Distinct antibody-mediated modifications of FLS resulting from phage display phenotypic screening

To use phage display phenotypic screening to identify novel components of FLS, phalloidin-stabilized structures were initially generated using PI(4,5)P_2_ supported lipid bilayers and *Xenopus* egg extracts and subsequently fixed with formaldehyde (Walrant et al., 2015). Since F-actin becomes densely polymerized within FLS, potentially restricting the access of phage capable of binding integral FLS components, screening was performed on three types of FLS structures: on early assemblies where FLS were fixed rapidly, on mature structures from later timepoints, and on a third preparation, in which mature structures were subjected to additional washing to relax the bundled FLS architecture. To enhance the likelihood of isolating antibodies targeting novel components, the phage library was deselected against a cocktail of purified actin regulatory proteins that are known to localise to FLS (actin, Toca-1, Ena, VASP, N-WASP, fascin and the Arp2/3 complex (Lee et al., 2010)), in addition to deselecting phage that bound the supported lipid bilayer and experimental set-up as depicted schematically in Figure 1A. Phage selected under each of the conditions (early, mature and washed FLS) were eluted, cloned, and sequenced. To confirm binding specificity, ELISAs were performed using phage identified uniquely from each of the three experimental conditions (Figure 1B). These were selected since, although the three conditions have antigens in common, phage binding under every condition are potentially more likely to engage residual proteins from the extracts and were consequently excluded from the initial analyses.

**Fig. 1.**
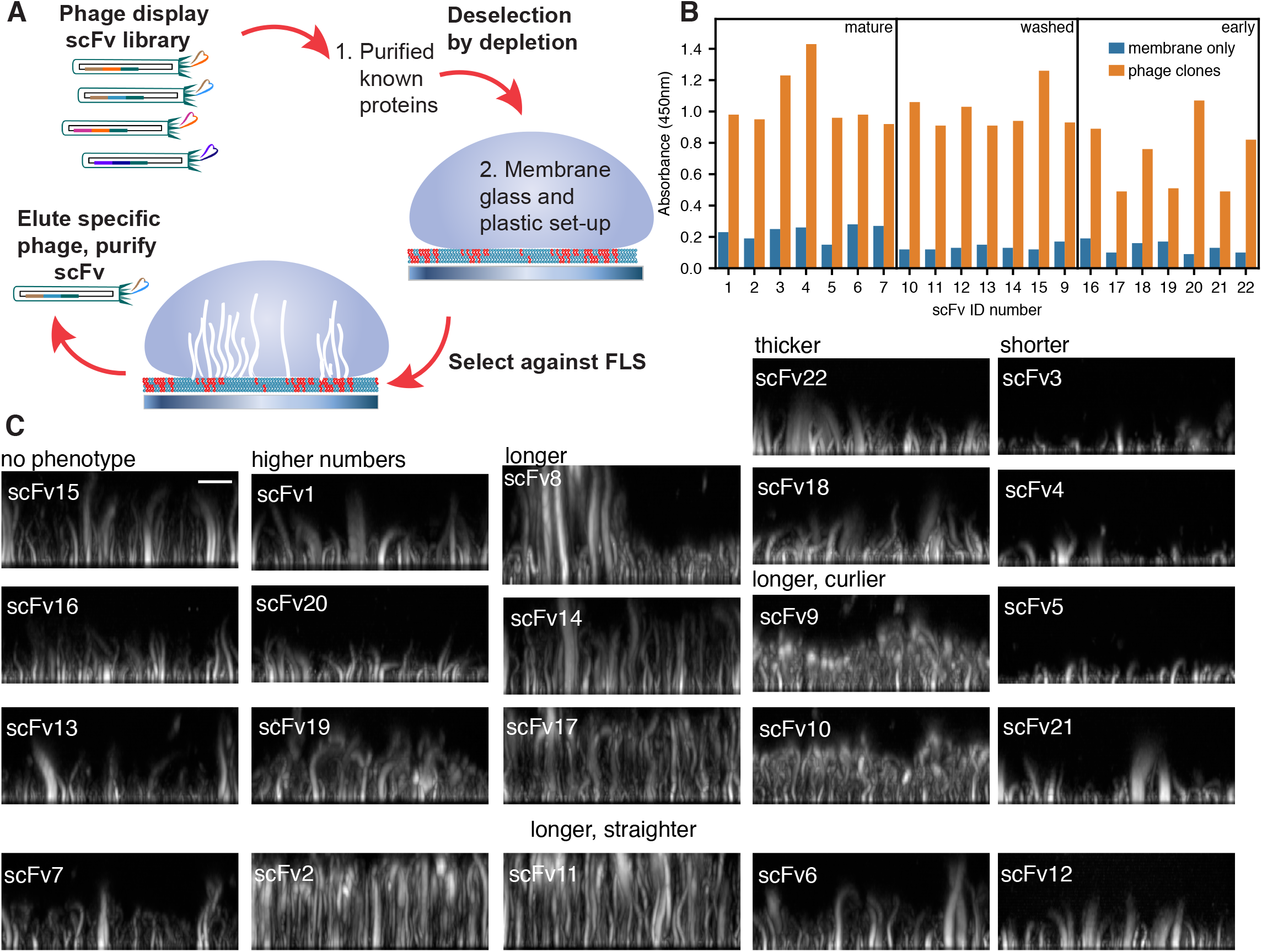
Phage display phenotypic screen on FLS. (A) Schematic of the scFv deselection and selection conditions (B) The specificity of scFv for different FLS was measured by ELISA against lipid bilayers, a mix of purified TOCA-1, fascin, Arp2/3 complex, N-WASP, Ena and VASP and mature FLS and only those with strong binding to FLS were selected. (C) FLS phenotypes on pre-addition of 5 μl of each scFv of ~ 1 mg/ml concentration to the FLS assay mix classified by the major phenotypes for each scFv, ranging from those that make FLS longer, curlier, shorter, straighter or thicker, or a mixture of more than one phenotype, quantified in Sup. Fig. 1. Images are maximum intensity projections of representative 1 μm confocal Z stack reconstructions viewed from the side (scale bar = 10 μm).

As a result of the screening and sequencing, 22 single chain variable region fragments (scFvs) were recombinantly expressed and purified to homogeneity. To investigate the potential effects of these scFvs on FLS, individual scFvs were pre-incubated with the assay mix including labelled actin and *Xenopus* egg extracts prior to the initiation of FLS formation upon addition to supported lipid bilayers mounted on glass coverslips. Emergent FLS were then visualised using spinning disk confocal microscopy.

Remarkably, a range of altered FLS phenotypes were evident upon co-incubation of different scFvs (Figure 1C), including changes in length (shorter, longer), morphology (thicker), stiffness (straighter, thicker) and increased number. We performed the screen with and without additional unlabelled actin as lengthening phenotypes could be constrained by limitations in the actin, whereas shortening phenotypes could be suppressed by greater actin concentrations. We quantified the morphological changes using our image analysis pipeline FLSAce (Dobramysl et al., 2019) (Figure S1A). As might be expected, changes in length were often associated with changes in stiffness (Figure 1C). Longer FLS could clearly be either straighter (scFv2) or curlier (scFv9), although this was not exclusively the case (*e.g.* scFv14 and scFv17). By contrast, induced thickening of the filopodia seemed to be an independent phenotype (Figure 1C*e.g.* scFv18 and scFv22). The phenotypes could also be more variable, *e.g.* scFv8 induced straighter FLS of both longer and shorter lengths (Fig. 1C, Sup. Fig. 1), which could be due to an increased persistence time of addition and loss of actin monomers. While we found some antibodies led to higher numbers of FLS, we have not yet identified antibodies that lead to fewer FLS, which may be because we pre-depleted the phage library against N-WASP and Arp2/3 complex which are needed for FLS initiation (Lee et al., 2010).This range of phenotypes suggested that the scFvs were potentially disrupting factors involved in different stages of FLS initiation, elongation and stabilization. Since the shorter phenotype could correspond to the inhibition or sequestration of factors involved in FLS formation and/or elongation, we focused our effort on the further characterisation of four scFvs (Figure 1C; scFv3, scFv4, scFv5 and scFv21), which following pre-incubation in the extracts generated significantly shorter FLS.

### SNX9 as a key mediator of FLS formation

To identify the target antigen/s of the four scFvs that generate shorter and straighter FLS, we initially used each scFv to probe *Xenopus* egg extracts by immunoblotting. All four scFvs recognised a common 70 kDa species, which was most evident with scFv3 and scFv4 (Figure 2A, Figure S2A). To enable subsequent immunoprecipitation, the two scFv variable regions with higher avidity were converted to an IgG1 framework to generate monoclonal antibodies (correspondingly, IgFLS3 and IgFLS4). These antibodies also recognised a common 70 kDa species when used to equivalently probe *Xenopus* egg extracts (Figure 2A; extract lanes, IgFLS3 and IgFLS4). Having confirmed their specificity, these antibodies were used to perform immunoprecipitations (IPs) from *Xenopus* egg extracts. Following separation by SDS-PAGE and size exclusion by excision of regions of the polyacrylamide gel spanning ~50-80 kDa, eluted proteins were identified by mass spectrometry by comparing extracts, IP with control IgG and IP with IgFLS3 (Figure S2B,C). Of 105 proteins unique to the IgFLS3 IP the highest scoring target of the right molecular weight was sorting nexin 9 (SNX9) (Figure S2D), a PX-BAR protein involved in endocytosis and intracellular trafficking. Indeed, purified SNAP-tagged SNX9 was also recognised by the monoclonal antibodies scFv3, scFv4 and scFv5 upon immunoblotting (Figure S2E).

**Fig. 2.**
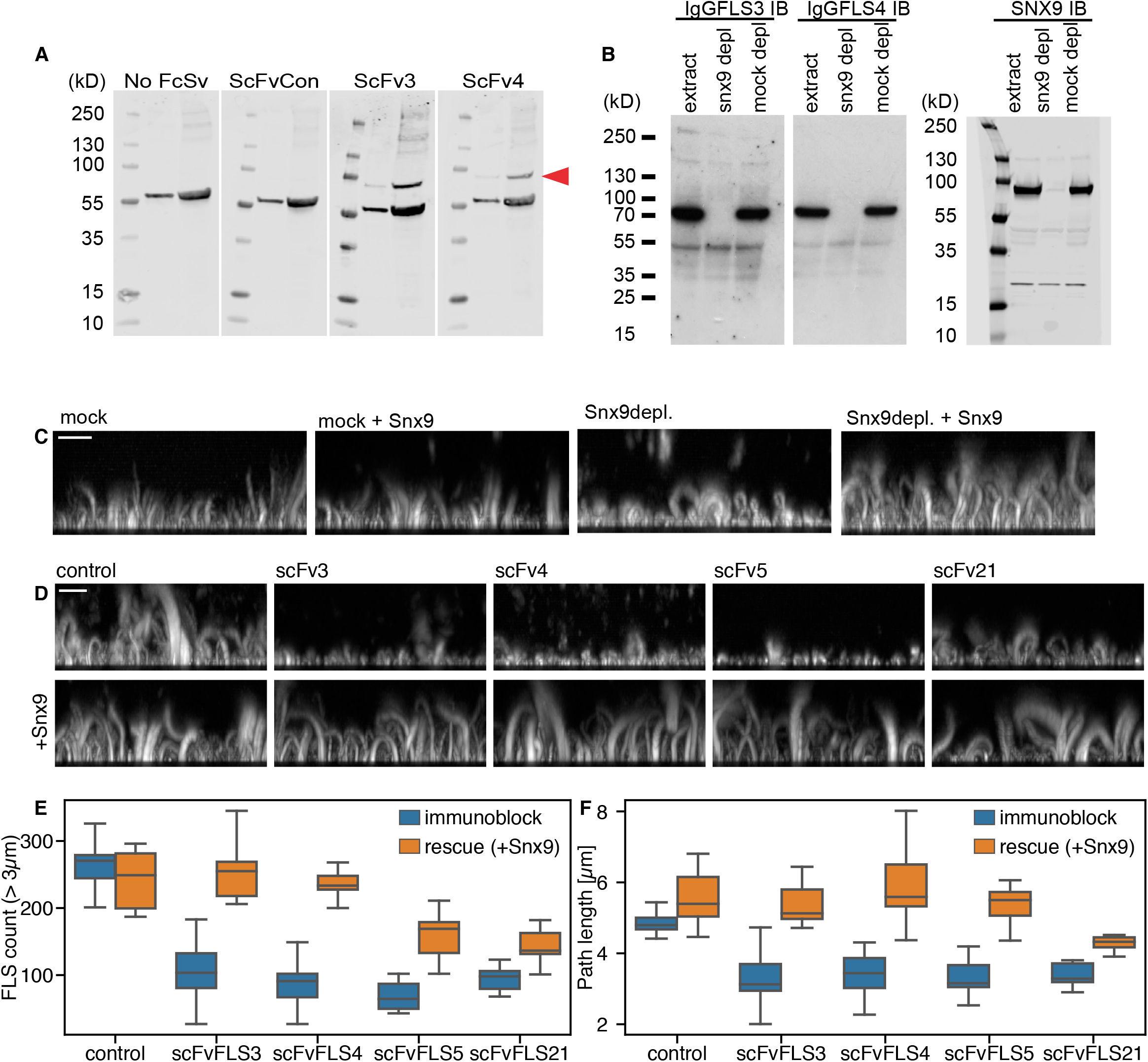
Identification of SNX9 as the antigen to antibodies giving a shorter FLS phenotype. (A) Western blot of two concentrations of *Xenopus* egg extracts (for each blot left lane = 25 μg, right = 100 μg extracts) using the indicated scFvs. ScFvFls3 and 4 both detect a specific band not detected in either control sample (red arrow). The band at ~60 kDa in all four blots is due to non-specific reactivity of the secondary or tertiary antibodies with extracts (as indicated by its presence in the no primary ScFv control). (B) Western blots of control extracts and SNX9 or mock immunodepleted extracts with either IgGFLS3, IgGFLS4 or rabbit-anti SNX9 antibody. In each case, the band at ~70kDa is specifically lost on SNX9 depletion. (C) Side maximum intensity projections of representative 1 μm confocal Z stacks of either mock or SNX9 immunodepleted extracts. Immunodepletion with anti-SNX9 antibodies reduces FLS length, which is rescued by addition of 20 nM purified SNX9 to the reaction mix (scale bar = 10 μm). (D-F) Immunoblock by specific scFvs and rescue by addition of 20 nM purified SNX9. Statistics are in Sup Table 1. (D) Side maximum intensity projections of representative 1 μm confocal Z stacks of FLS grown using extracts immunoblocked with control or specific scFVs, illustrating the reduction in FLS length and number, which can be rescued by addition of purified SNX9 (scale bar = 10 μm). (E-F) Quantification of the reduction in FLS number and length and rescue on SNX9 addition, showing the median, quartiles and range. Statistics are in Sup Table 1.

To validate the requirement for SNX9, we used a previously raised rabbit polyclonal antibody (Gallop et al., 2013) to immunodeplete SNX9 from *Xenopus* egg extracts. Depletion was assessed by immunoblotting the extracts with IgFLS3 and IgFLS4. This showed a complete loss of the previously recognised species at ~70 kDa (Figure 2B, compare ‘SNX9 depl’ and ‘mock depl’ samples on the IgFLS3 and IgFLS4 immunoblots, left). A comparable result was obtained when identical samples were alternatively immunoblotted with the anti-SNX9 antibody, definitively identifying the ~70 kDa species as SNX9 (Figure 2B, compare ‘SNX9 depl’ and ‘mock depl’ samples on the SNX9 immunoblot, right). In direct correlation with the immunoblock experiments (Figure 1C), FLS generated using the SNX9-depleted *Xenopus* egg extracts and PI(4,5)P_2_-supported lipid bilayers were significantly shorter (Figure 2C compare ‘mock’ and ‘SNX9 depl’ panels), a phenotype rescued by the addition of purified SNX9 (Figure 2B compare ‘mock+SNX9’ and ‘SNX9 depl + SNX9 panels). Furthermore, imaging and quantification revealed that the immunoblock induced by pre-incubation of extracts with scFvFLS3, scFvFLS4, scFvFLS5 or scFvFLS21 could be overcome by the addition of purified SNX9 (Figures 2D, 2E and 2F, statistics are in Sup Table 1). Although scFvFLS21 was less effective as an immunoblotting probe for *Xenopus* egg extracts (Figure S2C), scFvFLS21-dependent inhibition of FLS was similarly relieved following the addition of purified SNX9 (Figure 2C), suggesting that the antibody might recognize a conformational epitope present in the native protein rather than a linear epitope exposed upon denaturing SDS-PAGE or be bispecific for SNX9 and a binding partner. All together, these data identify SNX9 as an important mediator of FLS formation.

To establish directly whether SNX9 localizes to FLS, we supplemented *Xenopus* egg extracts with purified labelled SNAP-SNX9 and used PI(4,5)P_2_-supported lipid bilayers to generate FLS. Labelled SNX9 was clearly enriched at the tips of FLS proximal to the membrane from which the F-actin bundle emerged (Figure 3A, SNX9). SNX9 was co-incident with VASP, a marker for the tip complex of actin regulators present on the membrane (Figure 3A, VASP). Equivalent localisation was observed when fixed FLS were immunostained with an anti-SNX9 antibody (Figure 3B). In agreement with the immunoblock experiments, these data establish that endogenous or exogenously added SNX9 localises to the membrane-proximal tip, and is therefore likely a regulatory component rather than a structural element distributed throughout the FLS.

**Fig. 3.**
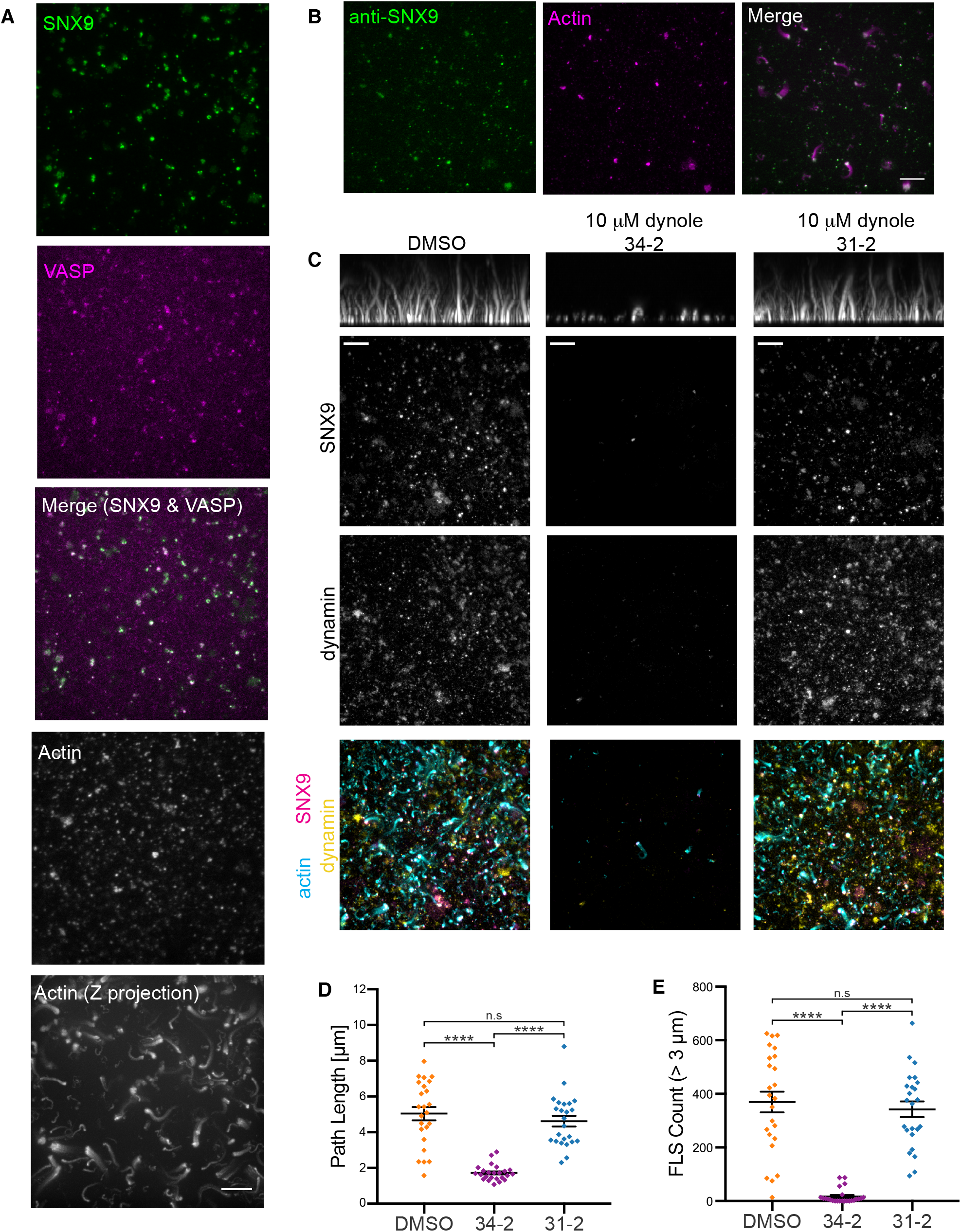
SNX9 and dynamin localise to FLS and FLS do not form in the presence of dynamin inhibitors. (A) TIRF imaging of a representative FLS region illustrating that purified SNX9 (green) and VASP (magenta) overlay with each other and actin (TIRF image labelled “actin” and Z-maximal projection of a 1 μm confocal stacks labelled “Actin (Z projection)”) at FLS structures (B) TIRF and confocal imaging of a representative FLS region immunolabelled using an anti-SNX9 antibody illustrating SNX9 (green) localised at FLS tips as marked by actin (magenta; TIRF in isolated channel, maximal projection of a 1 μm z-stack in merged image) (C) TIRF and confocal imaging of representative FLS regions containing labelled SNX9 (TIRF, magenta in merged image), dynamin-2 (TIRF, yellow in merged image) or actin (maximally projected confocal side or Z views, cyan in merged image) treated with either DMSO, or 10 μM of either the dynamin inhibitor dynole 34-2 or the inactive control compound dynole 31-2. SNX9 and dynamin both localize to FLS. This is lost on treatment with dynole 34-2, but not the inactive control compound, while similarly actin polymerisation is inhibited by the inhibitor but not the not inactive isomer. (D, E) Quantification of data in (C), illustrating the effect of dynamin inhibition on FLS length and number. Each datapoint represents an individual imaging region, lines indicate mean ± SEM. For each treatment N = 24 imaging regions. Statistical significance for both path length and FLS count (> 3 μm) wasassessed by ordinary one-way ANOVA with Holm-Sidak’s multiple comparisons test. Overall ANOVA for both path length (D) and FLS count (E) P < 0.0001 (****). For individual comparisons; in both (D & E) DMSO vs. dynole 34-2 and dynole 34-2 vs. dynole 31-2 P < 0.0001 (****), for (D) DMSO vs. dynole 31-2 P = 0.2901 (n.s.), in (E) DMSO vs. dynole 31-2 P = 0.4917 (n.s.). For all images, scale bar = 10 μm.

Dynamin is a key binding partner of SNX9, implicated in filopodia formation as well as in endocytosis (Chou et al., 2014). Given the location of SNX9, we next explored the potential relationship between dynamin and FLS formation, *Xenopus* egg extracts were supplemented with purified labelled-dynamin and SNX9 simultaneously, and PI(4,5)P_2_-supported lipid bilayers used to generate FLS as previously. Dynamin and SNX9 frequently co-localised at the membrane-proximal tips of FLS (Figure 3C, dynamin and SNX9). Indeed, the dynamin inhibitor dynole (34-2) but not its inactive isoform (31-2) blocked both dynamin and SNX9 recruitment to the PI(4,5)P_2_-supported lipid bilayers (Figure 3C), resulting in a significant quantifiable inhibition of actin polymerisation (Figure 3D) and FLS formation (Figure 3E). This reinforces the notion that a SNX9-dynamin containing complex located at the surface of the bilayer is an important mediator of FLS formation.

We have previously demonstrated synergistic interplay between SNX9, PI(4,5)P_2_ and PI(3)P underpinning actin polymerisation, defining SNX9 as a dual phospholipid sensor via binding of PI(4,5)P_2_ to the BAR domain and PI(3)P to the PX domain (Daste et al., 2017; Gallop et al., 2013). Since phosphoinositide 3-kinase (PI3K) is implicated in filopodia formation, through the generation of PI(3,4,5)P_3_ and PI(3,4)P_2_ (Jacquemet et al., 2019; Johnson and Haugh, 2016), we investigated the effect of inhibiting PI3K on FLS. Recruitment of SNX9 to PI(4,5)P_2_-supported lipid bilayers was markedly reduced in the presence of the PI3 kinase inhibitor LY294002, while the recruitment of an independent regulatory factor TOCA-1 remained unchanged (Figure 4A, compare SNX9 and TOCA-1). Again, a corresponding reduction in actin polymerisation and FLS formation was also evident (Figure 4A). Equivalent results were obtained using the general PI3K inhibitor wortmannin (compare PI(4,5)P_2_ only DMSO and wortmannin in Figure 4B to Figure 4A). Intriguingly, SNX9 recruitment and actin polymerisation could only be partially restored when the wortmannin-inhibited *Xenopus* egg extracts used with supported lipid bilayers were supplemented with exogenous PI(3)P but not those supplemented with PI(3,4)P_2_ or PI(3,4,5)P_3_, suggesting that at least *in vitro* PI(3)P, the terminal product of dephosphorylation of these more complex species, was the active constituent. Indeed, PI(3)P could be visualised using purified PI(3)P responsive mCh2xFYVE as a lipid reporter on the supported bilayers where FLS grow, and also upon restoration of SNX9 recruitment and actin polymerisation after wortmannin treatment (Figures 4B, 4C and 4D).

**Fig. 4.**
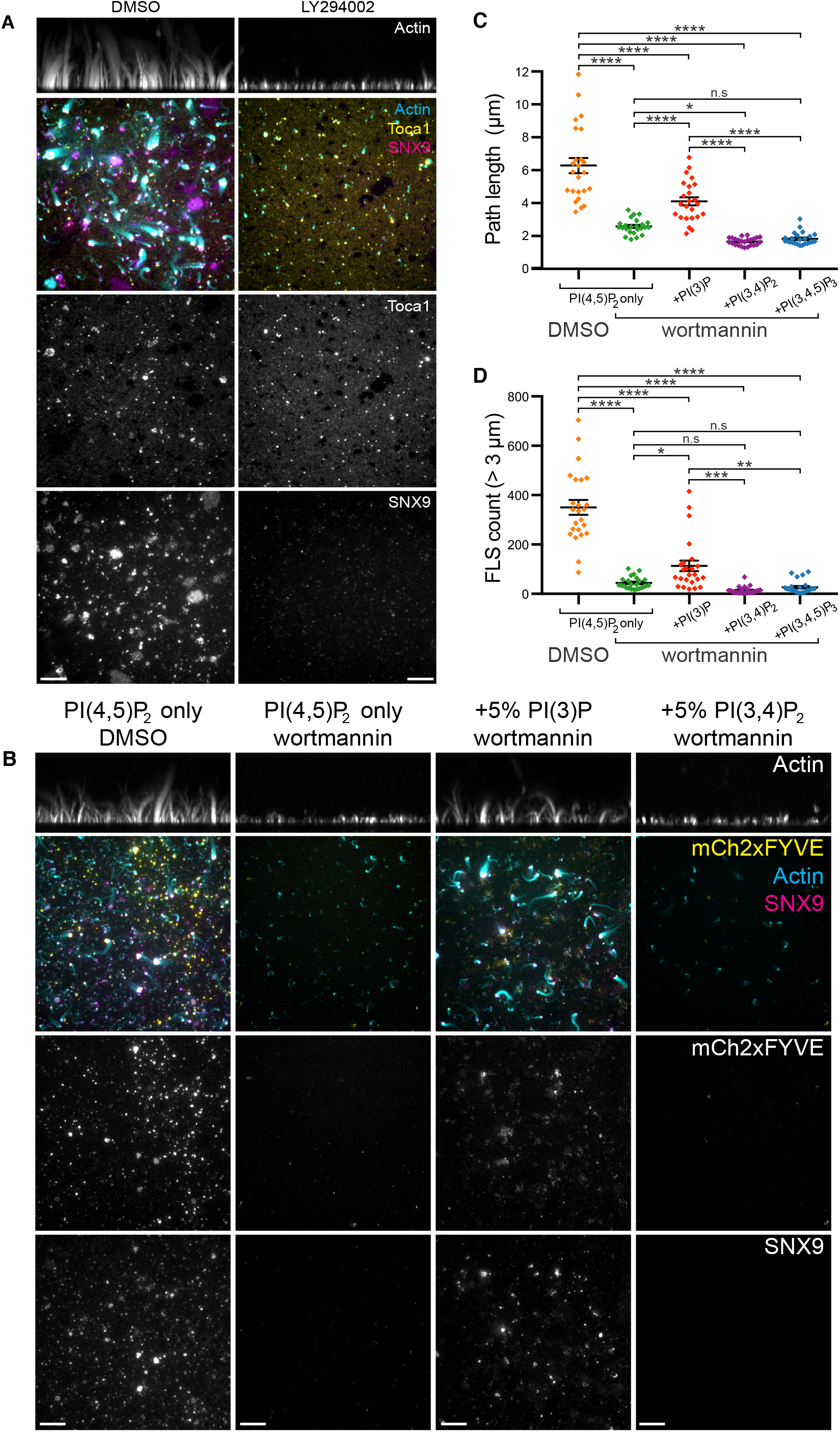
Class I PI 3-kinase is involved in FLS formation with PI(3)P being the key downstream lipid. (A) TIRF and confocal imaging of representative FLS regions containing labelled SNX9 (TIRF, magenta in merged images), TOCA-1 (TIRF, yellow in merged images) or actin (maximally projected confocal side or Z views, cyan in merged images) treated with either DMSO, or 100 μM of the general PI3K inhibitor LY294002. Both SNX9 presence and actin polymerisation at FLS is reduced on treatment with the inhibitor. TOCA-1 remains at the membrane and localises to the tips of residual actin structures. (B) TIRF and confocal imaging of representative FLS regions containing labelled SNX9 (TIRF, magenta in merged images), the PI(3)P probe mCh-2xFYVE (TIRF, yellow in merged images) or actin (maximally projected confocal side or Z views, cyan in merged images) grown on membranes containing either 10% PI(4,5)P_2_ or 10% PI(4,5)P_2_ supplemented with either 5% PI(3)P, or 5% PI(3,4)P_2_ and treated with either DMSO, or 2 μM of the general PI3K inhibitor wortmannin. Both SNX9 and mCh-2xFYVE localise to the membrane and FLS tips on treatment with DMSO; this is almost abolished on wortmannin treatment, coupled to loss of FLS actin structures. Inclusion of 5% PI(3)P in the supported lipid bilayer allows mCh-FYVE labelling to and SNX9 recruitment to the membrane to persist, and a partial rescue of actin FLS structures, while this does not happen when 5% PI(3,4)P_2_ is included instead. (C, D) Quantification of data in (B), illustrating the effect of PI3K inhibition by wortmannin on FLS length and number, and that this can be rescued by including PI(3)P but not PI(3,4)P_2_ or PI(3,4,5)P_3_ in the supported lipid bilayer. Each datapoint represents an individual imaging region, lines indicate mean ± SEM. For each treatment N = 24 imaging regions. Statistical significance for both path length and FLS count (>3 μm) was assessed by ordinary one-way ANOVA with Holm-Sidak’s multiple comparisons test. Overall ANOVA for both pathlength (C) and FLS count (D) P < 0.0001 (****). For individual comparisons; for all comparisons marked (****), P < 0.0001. In (C), PI(4,5)P_2_ + wort vs. +PI(3,4)P_2_ + wort, P=0.0247 (*) and PI(4,5)P_2_ + wort vs. +PI(3,4,5)P_3_ + wort, P =0.0583 (n.s). In (D),PI(4,5)P_2_ + wort vs. +PI(3)P + wort, P =0.0197 (*), +PI(3)P + wort vs. +PI(3,4)P_2_ + wort, P =0.0004 (***), +PI(3)P + wort vs. +PI(3,4,5)P_3_ + wort, P =0.0024 (**), PI(4,5)P_2_ + wort vs. +PI(3,4)P_2_ + wort, P =0.5066 (n.s) and PI(4,5)P_2_ + wort vs. +PI(3,4,5)P_3_ + wort, P=0.7199 (n.s). For all images, scale bar = 10 μm.

### SNX9 localizes to the tip and shaft of cellular filopodia

As exemplified by the previous experiments, the FLS system allows the identification of constituent components of filopodia and their biochemical analysis. It is also highly tractable, for instance allowing the isolation and manipulation of the structures themselves, the selective depletion of components from the extract, the application of pharmacological inhibitors, and the ability to add back both protein and lipid components to examine mechanistic dependency. While this allows for detailed analyses not possible in intact cells, there are of course disadvantages, for example the lack of a boundary membrane surrounding the filopodia, and the absence of integrated and contextual signalling pathways. It was therefore important to investigate whether SNX9 is a component of cellular filopodia in parallel.

Specialized filopodia called cytonemes play a role in transmitting Wnt signals in the dorsal marginal zone of *Xenopus* embryos which undergoes convergent extension movements to drive gastrulation of the embryo (Mattes et al., 2018). We dissected dorsal marginal zone explants from *Xenopus* embryos transiently co-expressing mCherry-SNX9 and the plasma membrane marker GFP-CAAX at stage 10.5. Live imaging revealed mCherry-SNX9 at the tips of extending cytonemes in this tissue, which could be simultaneously visualised using GFP-CAAX (Figure 5A, Figure S3A). This subset of SNX9-positive filopodia were frequently observed clustered together in distinct sub-regions of the explants, indicating these are potentially specialised structures (Figure 5A, Figure S3A, Sup movie 1).

**Fig. 5.**
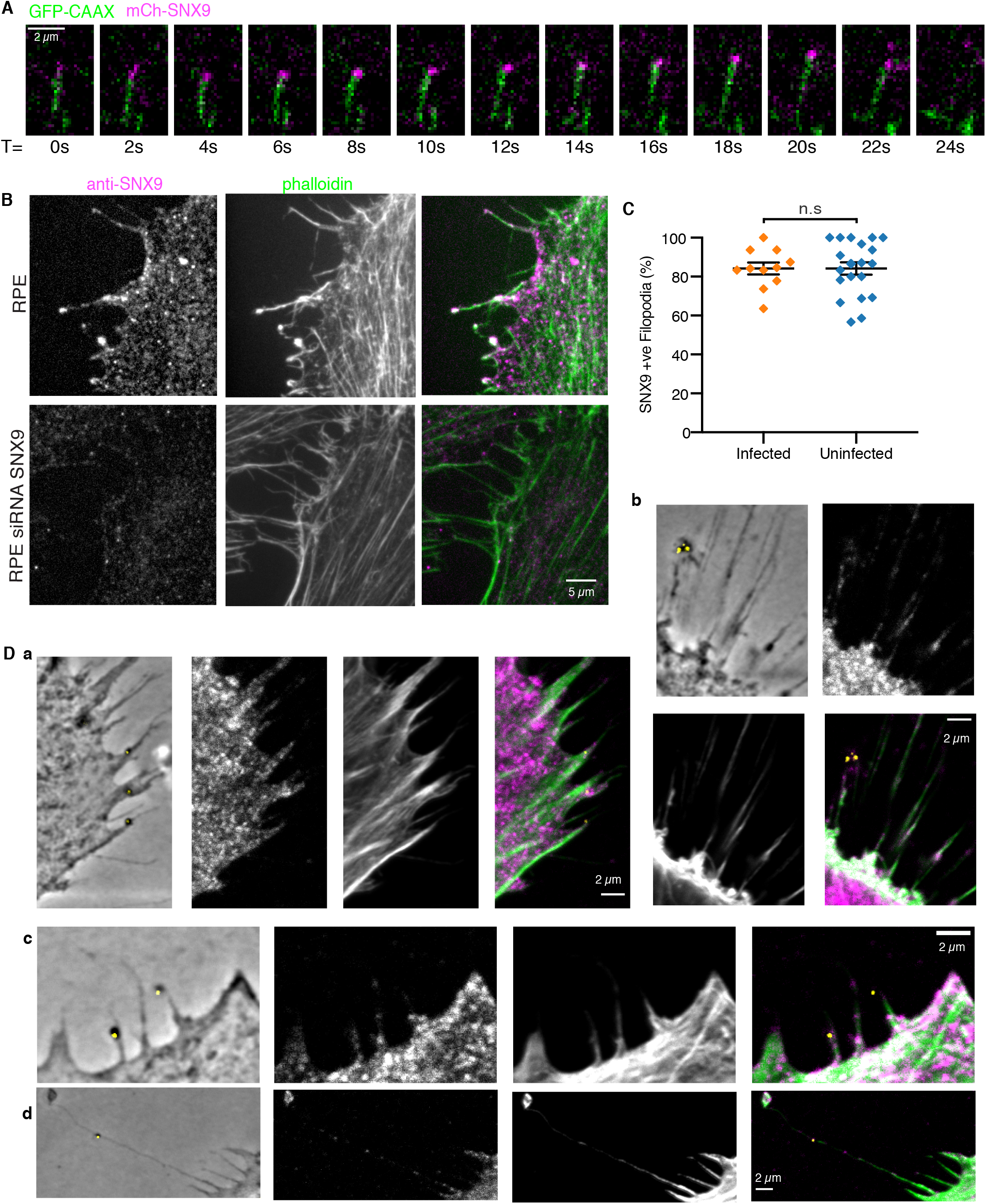
SNX9 localizes to filopodia in*Xenopus*gastrula explants and human cultured cells. (A) Background subtracted single frames from a TIRF microscopy timelapse sequence taken at 2 s intervals of *Xenopus* chordamesoderm cells expressing mCh-SNX9 (magenta) and GFP-CAAX membrane marker (green) showing SNX9 tracking a growing filopodium tip. Scale bar = 2 μm. (B) Confocal images of RPE cells treated with either scramble or SNX9 siRNA and immunolabelled with rabbit anti-human SNX9 (magenta) and phalloidin (green), illustrating presence of endogenous SNX9 in RPE cell filopodia and the abolition of SNX9 on siRNA mediated knockdown. Scale bar = 5 μm. (C) Quantification of SNX9 presence in filopodia in both uninfected RPE-1 cells and in cells infected with *C. trachomatis*. SNX9 immunolabelling stains ~85% of filopodia in untreated RPE-1 with no difference on *C. trachomatis* infection. Each data point represents an individual imaging region, lines indicate mean ± SEM. 248 filopodia from 62 cells across N=11 images were assessed for the infected condition, and 416 filopodia from 139 cells across N=20 images were assessed for the uninfected condition.Statistical significance was assessed by unpaired t-test, P = 0.9958 (n.s). (D a-d) Several example images of phase contrast and fluorescence confocal images of RPE cells infected with *C. trachomatis* (yellow) and immunolabelled for SNX9 (purple) and phalloidin (green), illustrating how *C. trachomatis* stick to pre-existing filopodia that contain SNX9 in either tip or shaft of the filopodia. Scale bars = 2 μm.

The endogenous localisation of SNX9 was investigated by indirect immunofluorescence of retinal pigment epithelial cells (RPE1), using two independent commercial human anti-SNX9 antisera. The specificity of these for SNX9 was confirmed as the immunofluorescence signal was markedly reduced following siRNA-mediated SNX9 knockdown (Figure 5B, Figure S3B (Ford et al., 2018)). Strikingly, SNX9 was present in ~90% of filopodia (Figures 5B and 5C). However, in contrast to the data in *Xenopus* embryos, the data from human cells suggest that SNX9 is a frequent component of filopodia, and that it can be present either in the tip or shaft.

SNX9 was recently reported to play a significant role during the induced entry of the obligate intracellular bacterium and major human pathogen *Chlamydia trachomatis* (Ford et al., 2018). Intriguingly, SNX9 was present at sites of early bacterial interaction with human cells and is involved in the filopodial capture of the extracellular pathogen. A correlative decrease in both filopodial number and bacterial internalisation was observed when SNX9^−/−^ knockout HAP1 cells were infected (Ford et al., 2018). This defect could be partially overcome by sedimenting the bacteria onto the cells, bypassing the SNX9-dependent capture step. Although SNX9 was present at entry sites and bacteria could be visualised in association with filopodia early during the encounter, the presence of SNX9 and its relationship to the adherent bacteria in these structures was not investigated in detail. Consequently, we examined the relationship between infectious *C. trachomatis* elementary bodies (EBs), endogenous SNX9 and filopodia during the initial minutes of host-pathogen interaction. As in the resting state, SNX9 was present equivalently at filopodia in infected RPE1 and HeLa cells (Figure 5C and S3, compare infected and uninfected quantification). *C. trachomatis* EBs were clearly observed in association with filopodia containing SNX9 (Figure 5D) and occasionally were also observed proximal to recruited SNX9 within the filopodial structure itself. Clusters or single infectious EBs were observed in association with the tips and shafts of SNX9-positive filopodia, which often appeared to capture EBs a considerable distance from the cell surface. Illustrative examples of these encounters are shown in Figure 5D panels a-d. These data capturing SNX9 within filopodia directly interacting with invading *C. trachomatis* EBs, significantly reinforce and extend the previous correlative data using SNX9^−/−^ knockout cells that implicated SNX9 as a cellular mediator of filopodia formation, a process subverted by a major human pathogen (Ford et al., 2018). Taken together, our data identify SNX9 as a major mediator of filopodial growth *in vitro*, in cultured human cells including in a known SNX9- and filopodia-dependent pathway subverted by a bacterial pathogen, and in the formation of specialised filopodia during convergent extension movements.

## Discussion

By performing a phage display phenotypic screen we identified antibodies against FLS that induce a variety of morphological changes in the resulting actin bundles. We show that this is a powerful approach for identifying novel proteins involved in the biogenesis of filopodia. By selecting the antibodies that generated shorter actin bundles, we identified the antigen recognised by four independent antibodies as SNX9 and subsequently demonstrated a functional role in FLS formation by immunodepletion and rescue experiments. Using human cell culture and *Xenopus* embryos explants we directly localized SNX9 to filopodia. The complexity of the phenotypes we obtained is reminiscent of the complex effects that individual regulatory proteins can have on the final morphology of actin within cells (e.g. the actin reorganisation prompted by cofilin-mediated disassembly). Here, the options include (1) an antibody blocking a single protein with a single activity, giving a distinct phenotype (2) blocking a single protein giving complex effects due to the multiple roles of the target protein or (3) the antibodies reacting with more than one sequence related protein and/or react against a protein complex giving either simple or complex phenotypes. As immunodepletion of SNX9 inhibits FLS formation similarly to the immunoblock and both immunoblock and depletion are alleviated with addition of SNX9, we can conclude that scFvFLS 3, 4 and 5 are blocking normal SNX9 function. Characterising the epitope on SNX9 that these antibodies bind will enable us to determine how SNX9 is contributing to filopodia formation and potentially the other cellular functions that use SNX9 (Bendris and Schmid, 2017). Previous work reveals some possibilities. In purified reconstitutions, SNX9 co-activates the Arp2/3 complex using its SH3 domain to bind N-WASP downstream of the PX-BAR domain interaction with PI(4,5)P_2_ and PI(3)P (Daste et al., 2017; Gallop et al., 2013; Yarar et al., 2007). In endocytosis, SNX9 interacts with dynamin via the SH3 domain and is implicated in membrane scission (Lundmark and Carlsson, 2004; Soulet et al., 2005), while dynamin itself has direct roles in polymerising actin (Gu et al., 2010; Lee and De Camilli, 2002). In cancer cells, SNX9 is thought to act by promoting Cdc42 activation but the protein domain responsible is not known (Bendris and Schmid, 2017; Bendris et al., 2016a; Bendris et al., 2016b). In FLS and filopodia formation, any or all of these mechanisms could apply. Here we show evidence implicating dynamin and PI(4,5)P_2_ and PI(3)P in FLS where Cdc42, N-WASP and Arp2/3 complex are also used (Lee et al., 2010).Arp2/3 complex activation has been associated with filopodia formation in some cases (Yang and Svitkina, 2011), while dynamin and Cdc42 also have recognized involvement (Chou et al., 2014).

In filopodia produced by expression of Myosin-X, enrichment of a membrane marker and localization of a PI(3,4)P_2_ probe has been observed at filopodia tips (Jacquemet et al., 2019), which could be related to our observations since SNX9 has been linked to PI(3,4)P_2_ as well as PI(4,5)P_2_ and PI(3)P (Daste et al., 2017; Gallop et al., 2013; Posor et al., 2013). How phosphoinositide conversion is dynamically regulated with filopodial protrusion and retraction and its integration with vesicle budding is another interesting avenue for future research.

We have shown that there is a SNX9-positive subset of filopodia in cells undergoing convergent extension movements during *Xenopus* gastrulation. These movements are known to be coordinated by Planar Cell Polarity (PCP)/noncanonical Wnt signalling (Heisenberg et al., 2000; Tada and Smith, 2000; Wallingford et al., 2000). PCP requires cell-cell communication mediated by asymmetric localisation of Wnt signalling proteins, where filopodia are proposed to play a role (Luz et al., 2014; Ossipova et al., 2015; Stanganello et al., 2015). In zebrafish neural plate patterning, Wnt8a puncta are observed on filopodia and filopodia are induced by PCP signalling via the tyrosine kinase receptor Ror2 (Luz et al., 2014; Mattes and Scholpp, 2018; Stanganello et al., 2015).SNX9 localisation to filopodia of *Xenopus* dorsal marginal zone cells during convergent extension may therefore point to a role for SNX9 in the transport of Wnt ligand to the signal receiving cell during PCP signalling. Wnt 8a puncta are released from filopodia onto receiving cells (Luz et al, PLoS One, 2014) and in early *Xenopus* embryo cleavage, vesicles are observed in association with filopodia (Danilchik et al., 2013; Luz et al., 2014). Because of its known roles in lipid binding, endocytosis and actin regulation, SNX9 is therefore a candidate molecule in linking membrane trafficking to filopodia formation.

Recently, SNX9 was shown to be important for the entry of *C. trachomatis* into mammalian cells (Ford et al., 2018). SNX9 localised to foci of early interaction, co-incident with the timepoint at which many infectious EBs are captured by filopodia.

Studies using SNX9^−/−^ knockout cells revealed a correlative defect in bacterial association with filopodia and cell entry rate. The data presented here now extend these studies. Rather than inducing SNX9 re-localisation for example by using a SNX9-binding translocated bacterial effector, our new findings suggest that *C. trachomatis* hijacks existing filopodia containing SNX9 to drive entry. While this does not preclude the existence of a SNX9-binding effector, it seems its role would be to anchor the adherent bacteria to SNX9-containing filopodia rather than to induce them. Indeed, while this entry mechanism shows some similarities with the determinants of FLS formation, for example the sequential generation of PI(3,4,5)P_3_ and PI(3)P from PI(4,5)P_2_ at entry sites (Ford et al., 2018), there are also intriguing differences. For instance, dynamin inhibitors do not interfere with chlamydial entry, but do inhibit the subsequent biogenesis of the nascent bacterial-containing vacuole and despite the presence of related phosphoinositide species, *C. trachomatis* entry is not sensitive either wortmannin or LY294002. Thus, as is typical of pathogen subversion of target cells, while this pathway undoubtedly shares some key hallmarks with FLS formation, it is likely that bacterial virulence factors replace or reprogram cellular factors to ensure deviation of bacterial-containing vacuoles from the canonical endocytic pathway.

To conclude, we report a new way of identifying proteins involved in filopodia and exploit this to identify a role for SNX9 in filopodia. This work has shed new light on the cellular mechanisms by which SNX9 is involved in development, cancer metastasis and pathogen entry and holds promise for further molecular dissection of FLS in understanding filopodia and actin regulation.

## Acknowledgements

We thank Charles Bradshaw for assembling databases and comparing *Xenopus* and mouse orthologues. Funding: This work was supported by European Research Council Grant 281971 and Wellcome Trust Research Career Development Fellowship WT095829AIA to JLG; a Medical Research Council Project Grant (N000846/1) to RDH. We used a Wellcome Trust grant 108467/Z/15/Z multi-user equipment grant funded mass spectrometer. We acknowledge the Gurdon Institute funding from the Wellcome Trust (092096) and CRUK (C6946/A14492). UD was supported by a Junior Interdisciplinary Fellowship Wellcome Trust grant No. 105602/Z/14/Z and a Herchel Smith Postdoctoral Fellowship. HS was supported by a Funai Foundation Overseas scholarship.

## Author contributions

Conceptualization, JLG and TJV; Methodology and Investigation, IKJ, JRG, AN, JM, HS, BM, CMM, LP, RDH, JLG; software and data curation UD and CMM, Writing - Original Draft, JLG, RDH, JRG, Writing - Review and Editing, CLD, JM, CMM, TJV, BM; Supervision, JLG, RDH, KSL, CLD, TJV; Project Administration, JLG, CLD, Funding acquisition, JLG.

## STAR Methods

### LEAD CONTACT AND MATERIALS AVAILABILITY

Further information and requests for resources and reagents should be directed to and will be fulfilled by the Lead Contact, Jennifer Gallop.Plasmids in this study will be deposited by Addgene on acceptance. Small quantities of the antibodies generated in this study are available on request.

### EXPERIMENTAL MODEL AND SUBJECT DETAILS

#### Cell lines

RPE-1 cells (RRID:CVCL_4388) are an immortalized line derived from human female retinal pigmental epithelial tissue. They were maintained in DMEM/F12 media (Sigma-Aldrich D6421) supplemented with 10% fetal bovine serum, 0.25% sodium bicarbonate (both Sigma-Aldrich), 100 μg/ml penicillin/100U/ml streptomycin and 2 mM L-Glutamine(all Gibco) at 37 °C in a humidified incubator containing 5% CO_2_, and were subcultured twice weekly.

#### Xenopus laevis

*Xenopus laevis* (RRID:XEP_Xla100) were used as both a source for the egg extracts used in FLS assays, and as a model organism in *Xenopus* embryo explant experiments.

Male and female *Xenopus laevis* were housed in the aquatic facility at the Gurdon Institute. *Xenopus* were housed in a Marine Biotech recirculating system, with 12 hour light/dark cycles and fed a commercial trout pellet diet. Female Xenopus laevis were used for egg collection only. To induce superovulation, adult female *Xenopus laevis* were injected with 150iU PMSG-Intervet (MSD Animal Health) 4 – 7 days pre-experiment and 400iU human chorionic gonadotrophin 12 – 18 hours pre-experiment. Primed *Xenopus* were kept in Marc’s modified Ringers’ (MMR) solution (100 mM NaCl, 5 mM Na-HEPES, pH 7.8, 2 mM KCl, 2 mM CaCl_2_, 1 mM MgCl_2_, 0.1 mM EDTA). Male *Xenopus laevis* were used for testes extraction. Male *Xenopus* were euthanised by subcutaneous injection of MS222. *In vitro* fertilization was performed by swirling mashed testis through collected eggs. The resultant embryos were injected with RNA at the two-cell stage, then maintained at 14°C, with explants taken at Nieuwkoop and Faber Stage 10. This research has been regulated under the Animals (Scientific Procedures) Act 1986 Amendment Regulations 2012 after ethical review by the University of Cambridge Animal Welfare and Ethical Review Body and covered by Home Office Project License P1B9A7D57 (License holder: Dr Jennifer L Gallop) and Home Office Personal licenses held by Jennifer L Gallop, Therese Jones-Green and Julia Mason.

## METHOD DETAILS

### Preparation of *Xenopus* egg extracts

High-speed supernatant *Xenopus* egg extracts were prepared according to our usual methods (Walrant et al., 2015). Briefly, eggs from up to 10 *Xenopus* were dejellied in MMR containing 2% w/v cysteine (adjusted to pH 8.0), then gently washed in *Xenopus* extracts buffer (XB: 100 mM KCl, 100 nM CaCl_2_, 1 mM MgCl_2_, 50 mM sucrose, 10 mM K-HEPES pH 7.4), followed by Extract buffer for CSF extracts (CSF-XB: 100 mM KCl, 10 mM K-HEPES, pH 7.4, 5 mM EGTA, 2 mM MgCl_2_, 50 mM sucrose). Following addition of protease inhibitor cocktail (Sigma-Aldrich P8340) and Energy Mix (final 1x concentration 2 mM ATP, 15 mM creatine phosphate, 2 mM MgCl_2_), eggs were transferred to thin-wall centrifuge tubes and crushed by centrifugation at 17,800*g* (in a SW-40 Ti rotor (Beckman-Coulter) for 10 mins at 4 °C. The crude cytoplasmic extract layer was extracted using a needle and diluted 10-fold in XB, then spun at 260,000*g* in a Ti-70 rotor (Beckman-Coulter) for 1 hour at 4 °C to prepare high speed supernatant. This supernatant was filtered through a 0.22μm syringe filter unit, then spin concentrated in a 10kDa MWCO spin concentrator (Amicon) to a final concentration of 25 mg/ml. Extracts were then supplemented with 200 mM sucrose and snap froze in liquid nitrogen for storage at −80 °C.

### Preparation of supported lipid bilayers

Supported lipid bilayers were prepared according to our usual methods (Walrant et al., 2015). Briefly, 2 mM total lipid liposomes were prepared containing, by mol fraction, 10% PI(4,5)P_2_, 30% phosphatidylserine (PS), and 60% phosphatidylcholine (PC) (Avanti Polar Lipids). For experiments which also included 5% PI(3)P, 5% PI(3,4)P_2_, or 5% PI(3,4,5)P_3_ (Avanti Polar Lipids) the amount of PC was reduced to 55%. To prepare liposomes, lipid stocks were mixed together and dried under a constant stream of nitrogen, then under vacuum for one hour, after which the lipid film was resuspended to the final 2 mM total lipid concentration in XB, and sonicated in a bath sonicator for 15 minutes. Liposomes were added to wells formed by silicone gaskets placed on glass coverslips or glass bottomed Proplate ®Microtiter Plates (Stratech Scientific) according to experiment and after adsorption and fusion to the glass surface washed several times with XB.

### Phage display screening & generation of antibodies

#### Isolation of anti-FLS antibodies

Supported lipid bilayers were prepared on Proplate ®Microtiter Plates. FLS assays were performed according to our usual methods (Walrant et al., 2015). Briefly, assay mix comprising 6 mg/ml *Xenopus* egg extracts, 2 mM DTT, 5 μM actin, energy regenerating system (Energy mix; final concentration 2 mM ATP, 15 mM creatine phosphate, 2 mM MgCl_2_) in XB was added to the supported bilayers and incubated for 15 min at room temperature (Walrant et al., 2015). For the mature FLS condition 2 μl phalloidin from a 10 mg/ml methanol stock was added and incubated for a further 5 min followed by 3 washes with XB and fixation with 4% formaldehyde for 75 min. For the washed FLS condition, after 15 mins FLS growth, 3 washes with XB were carried out then the FLS stabilised by addition of 2 μl phalloidin. After a 5 min incubation, assays were washed 3 times with XB and fixed with 4% formaldehyde in XB for 75 min. For the early FLS condition, the assay mix was removed after 3 min, washed 3 times with XB, 2 μl phalloidin added, incubated for 5 min then fixed with 4% formaldehyde in XB for 75 min. Fixed FLS were kept at 4 °C for up to 3 days. Phenotypic selections were carried out using the CAT2.0 human single-chain variable fragment (scFv) phage display library (Lloyd et al., 2009; Vaughan et al., 1996). Three rounds of selection were performed against each of three different filopodia structures: the mature, the washed and the early timepoint. Wells were washed with PBS and then blocked for 1 hour with PBS-Marvel milk powder (3% w/v). 10^9^−10^12^ phage were blocked for 1 hour at room temperature in PBS-Marvel milk powder (3% w/v) and then transferred to wells coated with lipid bilayers and irrelevant proteins to deplete non-specific binders. The blocked phage were subsequently incubated with the appropriate filopodia structure for 1 hour at room temperature and any unbound phage removed by a series of wash cycles using PBS-Tween (0.1% v/v) and PBS. Bound phage particles were eluted by addition of 5 μg/ml trypsin, infected into *E. coli* TG1 bacteria and rescued for the next round of selection (Vaughan et al., 1996).

#### Phage ELISA

Following three rounds of phage display selection, individual scFv were prepared as phage supernatants and screened by phage ELISA for binding to the corresponding filopodia structures used for selections (Osbourn et al., 1996; Vaughan et al., 1996). Nunc Maxisorp™ (Immobiliser™) plates (Thermo Fisher) coated with lipid bilayers and a mix of purified TOCA-1, fascin, Arp2/3 complex, N-WASP, Ena and VASP as described in (Dobramysl et al., 2019) were used for negative selection. Specific clones, defined as those which gave > 3-fold signal over background, were sequenced, expressed in TG1 Escherichia coli and purified via the C-terminal His tag by immobilized nickel chelate chromatography (Bannister et al., 2006; Lloyd et al., 2009)

#### Reformatting of scFv to IgG1 TM

Antibodies were converted from scFv to whole immunoglobulin G1 triple mutant (IgG1-TM, IgG1 Fc sequence incorporating mutations L234F, L235E and P331S) antibody format essentially as described by (Persic et al., 1997) with the following modifications. An OriP fragment was included in the expression vectors to facilitate use with CHO-transient cells and to allow episomal replication. The variable heavy (VH) domain was cloned into a vector containing the human heavy chain constant domains and regulatory elements to express whole IgG1-TM heavy chain in mammalian cells. Similarly, the variable light (VL) domain was cloned into a vector for the expression of the human light chain (lambda) constant domains and regulatory elements to express whole IgG light chain in mammalian cells. The plasmids were co-transfected into CHO-transient mammalian cells (Daramola et al., 2014) and IgG proteins purified from cell culture medium using Protein A chromatography. The purified IgG were analyzed for aggregation and degradation purity using SEC-HPLC and by SDS-PAGE.

### Cloning

All PCR oligonucleotide sequences can be found in the key resources table. *X. laevis* dynamin 2 (accession number BC142568) was PCR amplified from IMAGE clone 7979408 (Source Bioscience), and cloned into pCS2 his SNAP acceptor vector. *X. laevis* SNX 9 (accession number BC077183) was PCR amplified from IMAGE clone 3402622 (Source Bioscience) and cloned into either pCS2 Nterm GFP or pCS2 Nterm mCherry. GFP-Utrophin-CH was PCR amplified from a pCS2 construct (gift from Sarah Woolner, (Burkel et al., 2007)) and cloned into pGEX FA acceptor vector. All other constructs used have been previously described; see key resources table for details.

### Protein purification

All chemicals and reagents were from Sigma-Aldrich, unless otherwise stated. 6xHis-SNAP-TOCA-1 and 6xHis-GFP-Utrophin in pET or pGEX plasmids respectively were transformed into BL21 pLysS *E.coli*, with protein expression induced overnight at 19 °C. 6xHis-SNAP-SNX9 and 6xHis-SNAP-Dynamin2 in pCS2 constructs were transfected into 293F cells by 293fectin reagent (Thermo-Fisher Scientific) according to manufacturer’s instructions, and were harvested 48 hours later. All purification steps were performed at 4°C. Bacteria or 293F cells were harvested by centrifugation and resuspended in a buffer containing 150 mM NaCl, 20 mM Na-HEPES (pH 7,4), 2 mM 2-mercaptoethanol and EDTA-free cOmplete protease inhibitor tablets, after which they were lysed by probe sonication. Lysates were spun at 40,000 rpm for 45 mins in a 70Ti rotor (Beckman-Coulter), with supernatants then applied to Ni-NTA agarose beads (Qiagen) for affinity purification. Proteins were eluted by applying stepwise increasing concentrations of imidazole (50-300 mM) in a buffer containing 150 mM NaCl, 20 mM Na-HEPES (pH 7,4), 2 mM 2-mercaptoethanol. 6xHis-KCK-VASP in the pET15b plasmid was transformed into Rosetta DE3 pLysS cells and expression induced as for the pET constructs above. The protein was purified as above, with the following differences; all buffers contained 300 mM NaCl rather than 150 mM, and the beads used for affinity purification were cobalt agarose beads (Talon superflow, GE healthcare). For all proteins, fractions containing eluted protein were further purified using S200 gel filtration on an AKTA FPLC (GE healthcare) in a buffer containing 150 mM NaCl (300 mM for KCK-VASP), 20 mM Na-HEPES pH7.4, 2 mM EDTA and 5 mM DTT. Purification was verified by SDS-PAGE electrophoresis and Coomassie staining, with positive fractions pooled and concentrated in a 10 kDa MWCO spin concentrator (Millipore). Concentration was verified by A280 measurements on a Nanodrop, and proteins had 10% glycerol added before they were snap frozen in liquid nitrogen and stored at −80 °C.

#### Chemical labelling of proteins with fluorescent dyes

SNAP-tagged TOCA-1, SNX9 and Dynamin-2 were labelled using SNAP-Surface Alex Fluor 488 or Alexa Fluor 647 (New England Biolabs). 5-10 μM final concentration of protein was mixed with 10 μM SNAP-dye in a buffer containing 150 mM NaCl, 20 mM Na-HEPES (pH 7.4), 1 mM DTT and 1% (v/v) TWEEN 20 and incubated under gently rotation at 4 °C overnight. The labelled protein was dialysed into a buffer containing 150 mM NaCl, 20mM Na-HEPES (pH 7.4) and 10% glycerol in a 0.1 ml 20 kDa MWCO Side-A-Lyzer MINI Dialysis device (Thermo Fisher) to remove excess dye; dialysis occurred in two rounds over a 24 hour period. KCK-VASP was labelled at 4o C overnight by adding a 10-20 fold molar excess of the Alexa Fluor 568 maleimide dye (Thermo Fisher) to a 50 μM final concentration of the protein in the presence of a 10-fold molar excess of TCEP under gentle rotation. Excess dye was removed by buffer exchange into 300 mM NaCl, 20 mM Na-HEPES (pH 7.4), 10% glycerol using an Ultra-15 centrifugal filter unit spin concentrator (Amicon) with an Ultracel-10 membrane (Millipore).

### Immunodepletion

SNX9 was immunodepleted from *Xenopus* egg extracts using a Rabbit anti-*Xenopus* SNX9 antibody. Three rounds of depletion were performed using Protein A beads. Briefly, 300 μl antibody serum (SNX9 and pre-bleed serum as mock sample) was bound to 75 μl Dynabead protein A beads for 1 hour at room temperature with continuous rotation. Beads were washed with PBST, PBST plus 500 mM NaCl and XB buffer and then divided into three 25 μl aliquots. 100 μl *Xenopus* egg extracts were incubated subsequently with each aliquot of beads for 30 minutes at 4°C with rotation for each incubation. The resulting depleted and mock depleted egg extracts were analysed by Western blotting with IgFls3, IgFls4 and Rabbit anti-*Xenopus* SNX9 antibody utilized as primary antibodies, and Mouse anti-Human IgG1 Fc Secondary Antibody, HRP secondary antibody for the IgGs, and IRDye 800CW goat anti-rabbit IgG secondary antibody for the SNX9 antibody.

### FLS assays

FLS assays were performed according to our usual methods on glass coverslips in wells defined by silicone gaskets (Walrant et al., 2015). The basic reaction mix was composed of a 1:6 dilution of 25 mg/ml *Xenopus* egg extracts, 2 mM DTT, energy mix, 1 μM unlabeled rabbit skeletal muscle actin (Cytoskeleton), and 210 nM fluorescently labelled actin (either Alexa 488, 568 or 647, Thermo Fisher), made up in XB, and supplemented with other labelled proteins, antibodies or inhibitors as necessary the specific experiment. 50 μl of reaction mix was gently added to the supported lipid bilayer and incubated at room temperature for 25 mins before imaging. For the screening of the scFvs, 5 μl of the individual scFv was added to the basic reaction mix. For experiments involving the rescue of FLS prepared using mock/SNX9 immunodepleted or scFV treated extracts, 20 nM of SNX9 was added, and rather than using fluorescently labelled actin, 33 μM of GFP-utrophin was used to label FLS (Burkel et al., 2007). For experiments in which the localization of fluorescently labelled recombinant purified proteins was analysed, they were included in the reaction mix at the following concentrations; 20 nM VASP, 30 nM SNX9, 25 nM dynamin, 10 nM TOCA-1, 1 μg/ml mCh-2xFYVE. For experiments involving the addition of inhibitors to FLS, inhibitors (or appropriate dilutions of the vehicle, DMSO) were prepared as 10x stocks in XB, with 5 μl then added to complete the 50 μl reaction mix. The mixes were then incubated for 10 minutes before being added to the supported lipid bilayer to start the experiment. Inhibitors were used at the following final concentrations; 10 μM dynole 31-2, 10 μM dynole 34-2, 2 μM wortmannin, 100 μM LY294002.

### Immunolabelling of FLS

FLS were prepared using an assay mix comprised of 6 mg/ml *Xenopus* egg extracts, 2 mM DTT, Energy mix, 210 nM fluorescently Alexa Fluor 568 labelled actin made up in XB was added to the supported bilayers and incubated for 15 min at room temperature after which 0.5 μl phalloidin from a 10 mg/ml methanol stock was added. Following a further 5 min incubation, FLS were washed twice in XB then fixed in 4% formaldehyde made up in Cytoskeletal buffer (CB; 10 mM MES, 138 mM KCl, 3 mM MgCl_2_ and 2 mM EGTA) for 1 hour. Following 3x PBS washes, FLS were blocked in 2% BSA-PBS for 1 hour. Primary antibody mixes were made up in PBS; FLS were stained with the rabbit anti-*Xenopus* SNX9 antibody (1:100 dilution) for 1 hour. Following 3x PBS washes, the secondary antibody mix (goat anti-rabbit Alexa Flour 488 made up in PBS) was added for 1 hour. Fixed and labelled FLS were washed three further times in PBS then immediately imaged.

### Western blotting using FLS ScFvs and IgG1-TMs

Samples of either *Xenopus* egg extracts or purified recombinant SNX9 were run on 4 – 20 % mini-PROTEAN precast protein gels (Biorad) and transferred to nitrocellulose membrane using an iblot 2 dry blotting system. Western blots using ScFvs were blocked in 10% milk/PBST (1 x PBS + 0.1% Tween), incubated with 4.5 μg/ml ScFv in 1% milk/PBST for 1 hour at room temperature, washed with PBST, and then incubated overnight with a 1/1000 dilution in 1% milk/PBST of either rabbit anti-his antibody (Abcam) or mouse anti-myc antibody (Roche). After washing the membranes, tertiary incubations were performed with a 1/10,000 dilution in 1% milk/PBST of either anti-rabbit or anti-mouse 800 CW antibody (LI-COR Biosciences), washed, and imaged using an Odyssey Sa Reader (LI-COR Biosciences). Western blots using Fls IgGs were blocked in 5% milk/TBST (1 x TBS + 0.1% Tween). Antibody incubations were performed in antibody diluted in 5% milk/TBST. Fls IgGs were diluted to 2.5 μg/ml and the secondary mouse anti-Human IgG1-HRP antibody (Thermo Fisher) used at 1/500 dilution. HRP was detected using the ECL Prime western blotting system (GE Healthcare).

### Immunoprecipitation

For immuno-precipitation of IgG-FLS antigens, we employed a gravity flow column format as follows, with all steps conducted at 4°C. 2 ml of Affigel® 10 matrix (BioRad) slurry (1 ml column bed volume) was added to a poly prep chromatography column (BioRad) and washed with 3 bed volumes of ice cold water under gravity flow. IgG antibody was incubated with the immunoaffinity resin at a concentration of 1 mg/ml in 20 mM HEPES pH 7.4; 1 x PBS and columns were rotated overnight at 4°C. Affigel® activated affinity resin binds to free primary amines and therefore it was necessary to block unreacted residues in order to avoid non-specific protein binding, by the addition of 100 μl of 1 M ethanolamine (pH8.0) spiked into the resin/antibody mix. The column was left to rotate for 1 hour at 4°C. Unbound supernatant was eluted from the resin under gravity flow and the column was washed with 8 column bed volumes of 20 mM HEPES pH 7.4. The efficiency of antibody binding to the matrix was evaluated by SDS-PAGE before proceeding to the next step. 1 ml of *Xenopus* egg extracts (approx. 12 mg/ml concentration) were mixed with 20 μl 10 % Tween, 200 μl 10x XB and 780 μl water and loaded onto the IgG-bound column. The columns were rotated overnight at 4°C in the presence of the antigen-containing extracts. Unbound supernatant was removed from the column under gravity flow and the column was washed with 10 ml 0.5 M NaCl, followed by three 10 ml washes with binding buffer (20 mM HEPES pH 7.4, 1 x PBS). All flow through and wash samples were retained for further analysis. Elution of the antibody-bound antigen was enabled by ten sequential additions of 250 μl elution buffer (200 mM Glycine pH 2.5, 150 mM NaCl). Each of the ten eluates were collected into separate tubes and immediately neutralised by the addition of 50 μl 1 M Tris pH 8.0. In total, three samples were processed in parallel: (i) immuno-precipitation with the FLS-IgG of interest, (ii) immunoprecipitation with a control IgG antibody was used to assess non-specific binding to IgG, and (iii) a column with resin was prepared (as above) and incubated with extracts without the addition of IgG, to account for non-specific binding of proteins to the resin matrix.

### Proteolytic digestion and LC-MS/MS mass spectrometry

Immunoprecipitated eluates from (i) IgFls3 IgG, (ii) control IgG and (iii) *Xenopus* egg extracts + resin, were concentrated by vacuum centrifugation (Labconco), run on 4–15% Mini-Protean TGX gels (Bio-Rad) and stained with Instant Blue™ coomassie stain (Sigma Aldrich). The fourth elution of the ten sequential eluates was processed for liquid chromatography tandem mass spectrometry (LC-MS/MS), as this was the sample containing the antigen of interest identified by immunoblotting. Bands were excised from the coomassie gel corresponding to the 50 – 80 kDa region and processed for mass spectrometry as follows. Gel pieces were cut into 1-2 mm cubes, destained by several washes in 50 % (v/v) acetonitrile, 100 mM ammonium bicarbonate solution and then dehydrated with 100 % (v/v) acetonitrile. Disulphide reduction was achieved by incubation with 10 mM dithiothreitol (DTT) at 37°C for 1 hour, followed by alkylation of cysteine residues with 5 mM iodoacetamide at room temperature protected from light for 1 hour, where both solutions were made in 100 mM ammonium bicarbonate. Gel pieces were washed several times in 50 % (v/v) acetonitrile, 100 mM ammonium bicarbonate solution at 37°C, before being dehydrated in 100 % (v/v) acetonitrile. Proteolytic digestion was achieved by the addition of 50 μl sequencing-grade modified trypsin (Promega) at a concentration of 10 ng/μl in 50 mM ammonium bicarbonate. A further aliquot of trypsin solution was added after one hour to ensure gel pieces remained hydrated during overnight incubation at 37 °C. The supernatant containing the eluted peptides was retained and analysed by LC-MS/MS analysis with a Waters nanoAcquity UPLC (Thermo Fisher Scientific) system coupled to an Orbitrap Velos mass spectrometer (Thermo Fisher Scientific). Peptides were first loaded onto a pre-column (Waters UPLC Trap Symmetry C18, 180-μm i.d. × 20 mm, 5-μm particle size) and eluted onto a C18 reverse-phase column at a flow rate of 300 nl / min (nanoAcquity UPLC BEH C18; 75-μm i.d. × 250 mm, 1.7-μm particle size, Waters, UK) using a linear gradient of 3 – 40 % Buffer B over 40 minutes (60 min total run time including high organic wash and re-equilibrium steps). Buffer A: 0.1 % formic acid in HPLC-grade water (v/v); Buffer B: 0.1 % formic acid in acetonitrile (v/v). The mass spectrometer was operated using data dependent MS/MS acquisition in positive ion mode. Full scans (380 – 1500 m/z) were performed in the Orbitrap with nominal resolution of 30,000. The top 20 most intense monoisotopic ions from each full scan were selected for collision induced dissociation (CID) in the linear ion trap with a 2.0 m/z precursor ion selection window and normalized collision energy of 30 %. Singly charged ions were excluded from MS/MS and a dynamic exclusion window of ± 8 ppm for 60 sec was applied.

#### *Xenopus* embryo explants

##### Capped RNA synthesis

CAAX-GFP and mCherry-SNX9 capped RNA was synthesized using the mMESSAGE mMACHINE SP6 transcription kit (Invitrogen) from NotI linearized pCS2 CAAX-GFP and pCS2 his mcherry SNX9 plasmids. Linear DNA was cleaned up prior to transcription using the QIAquick PCR purification kit (Qiagen). 1 μg purified linear DNA was added to a standard mMESSAGE mMACHINE transcription reaction with the addition of 2 U/μl of Murine RNase inhibitor (New England Biolabs) and incubated for 2 hours at 37 °C. Unincorporated nucleotides were removed using the RNeasy mini kit (Qiagen).

##### RNA injection and Keller explants

9.2 nl of a solution containing both capped RNAs at 30 ng/μl was injected into each cell of dejellied 2 cell *Xenopus* embryos maintained in MMR containing 4% Ficoll. The injected embryos were transferred to 0.1 x MMR solution after undergoing a minimum of one cell division and cultured at 14 °C. Briefly, to visualise *Xenopus* chordamesoderm cells, explants of dorsal mesendoderm and ectoderm were prepared from the injected embryos at the early gastrula stage (Keller and Danilchik, 1988). The vitelline membrane was removed and a section of dorsal mesendoderm and ectoderm extending from the bottle cells removed using an eyebrow knife and hair loop and cleaned of loose mesoderm cells. Explants were placed under a glass bridge in DFA buffer (53 mM NaCl, 32 mM Na-gluconate, 4.5 mM K-gluconate, 1 mM CaCl_2_, 1 mM MgSO_4_, 5 mM Na_2_CO_3_, pH 8.3 with HEPES) for 3 hours at room temperature, and then imaged.

### Immunostaining of cells and *Chlamydia* infection

RPE1 cells were infected with wild-type *C. trachomatis* LGV2 as described previously (Ford et al., 2018). Briefly, cells were cultured on coverslips and infected with *C. trachomatis* at a multiplicity of infection of 5-30. To synchronise *Chlamydia* entry, bacteria were spinoculated at 900*g* for 10 min. After 30 minutes at 37°C, samples were fixed and immunostained. For staining with mouse anti-human SNX9 antibody samples were fixed in 4% (w/v) paraformaldehyde in phosphate buffered saline (PBS), for 20 minutes at room temperature. After fixation, samples were blocked in blocking solution (10% fetal calf serum (FCS) in PBS + 0.2% (w/v) saponin) for 1 hour, and then incubated for 1h with primary antibody (1/100 in PBS + 3% FCS + 0.2% saponin), washed 3 times in PBS, and finally incubated for 1 hour with goat anti-mouse IgG coupled to Alexa Fluor 568 (1/500 in PBS + 3%FCS + 0.2% saponin), phalloidin coupled to Alexa Fluor 488 (Thermo Fisher, 1/100), and DAPI (Thermo Fisher, 1μg/mL). For staining with rabbit anti-SNX9 antibody, samples were fixed according to the ‘Golgi fixation method’ described by Hammond and co-workers (Hammond et al., 2009). Samples were fixed for 15 minutes in 2% (w/v) formaldehyde in PBS. After fixation, samples were permeabilized for 5 minutes in permeabilization solution (20 μM digitonin, in buffer A: 150 mM NaCl, 20 mM HEPES, and 2 mM EDTA). Samples were blocked in blocking solution (10% goat serum in a buffer A) for 45 minutes, prior to incubation for 1h with primary antibody (diluted 1:100 in buffer A + 1% goat serum), and then for 45 minutes with goat anti-rabbit IgG coupled to Alexa Fluor 647 (1/1000 in buffer A + 1% goat serum), phalloidin coupled to Alexa Fluor 488, and DAPI. Samples were finally fixed post staining for 5 minutes in 2% (w/v) formaldehyde solution.

### siRNA treatment

siRNA (see Key Resources Table for sequences) was used to knockdown SNX9 expression in RPE-1 cells. Cells were seeded at an appropriate density on glass coverslips or tissue culture plates. 5 μl anti-SNX9 siRNA (custom sequence as used in (Nandez et al., 2014), synthesized by Dharmacon) or non-targeting control (Dharmacon) siRNA was added to 250 μl OptiMEM, with 8 μl Lipofectamine-3000 mixed with a second 250 μl OptiMEM aliquot. After five minutes these were combined, and after 20 minutes the siRNA complexes added to the cells; 200 μl was added to a six-well plate well, 50 μl to a 24-well plate well. A two shot protocol was used in which cells were transfected twice at both 24 and 72 hours after seeding, with analysis (immunolabelling or western blotting) occurring 24 hours after the second shot.

### Microscopy

Imaging of all FLS experiments and live *Xenopus* explant samples were performed on a custom combined spinning disk/total internal reflection (TIRF) fluorescence microscope supplied by Cairn research. The system was based on an Eclipse Ti-E inverted microscope (Nikon), fitted with an X-Light Nipkow spinning disk (CREST), an iLas2 illuminator (Roper Scientific), a Spectra X LED illuminator (Lumencor) and a 250 μm piezo driven Z-stage/controller (NanoScanZ). Images were collected at room temperature through a 100x 1.49NA oil objective using a Photometrics Evolve Delta EMCCD camera in 16-bit depth using Metamorph software (Version 7.8.2.0, Molecular devices). Alexa Fluor 488 and GFP samples were visualized using 470/40 excitation and 525/50 emission filters, Alexa Fluor 568 and mCherry samples with 560/25 excitation and 585/50 emission filters, and Alexa Fluor 647 samples with 628/40 excitation and 700/75 emission filters. For FLS assays, specific proteins of interest at the FLS tip (at the membrane) were imaged in TIRF, in conjunction with a confocal z-stack of the actin structure. Live time-lapse *Xenopus* explant samples were imaged in TIRF, with images captured every 2s. Cell culture experiments were imaged using a Zeiss LSM 710 laser scanning confocal microscope. All images were processed in FIJI (ImageJ (Schindelin et al., 2012)). Background subtracted images of *Xenopus* explant time-lapse images were prepared by subtracting the average intensity of the entire time-course from each individual timepoint.

## QUANTIFICATION AND STATISTICAL ANALYSIS

Data related to FLS count and FLS physical parameters was extracted using our Fiji (Image J) image analysis plugin FLS Ace, described in detail in (Dobramysl et al., 2019). Briefly, FLS structures are segmented by applying a 2D Difference of Gaussians filter with subsequent thresholding to each z-slice. FLS positions are then traced through the stack starting at the base image by employing a greedy algorithm. From this, the FLS morphology such as length, base area and straighness are extracted.

Peptide and protein identification was conducted with the Proteome Discoverer platform (version 1.4, Thermo Fisher Scientific) using the Mascot search algorithm (version 2.6, Matrix Science, London, UK) and searched against a database which combined Xenbase 9.1 and a database obtained from (Wuhr et al., 2014) (XL_9.1_v1.8.3.2.Wuhr_et_al_2014_Combined_cdhit95.fasta) containing 75,603 sequence entries, with the *Xenopus* accession numbers annotated by comparison to mouse orthologs. Fixed modification of carbamidomethyl (C) and variable modifications of oxidation (M) and deamidation (NQ) were selected. Protein grouping was enabled, minimum ion score of 20 was required and strict parsimony rule was applied. Trypsin was selected as the enzyme of choice and up to 2 missed cleavages were accepted. Precursor mass tolerance of 25 ppm and a fragment mass tolerance of 0.8 Da was used. Percolator (version 1.17) validation was applied using a high confidence FDR threshold of 0.01 based on q-value. Additional filters included a minimum protein score of 50 and a requirement of 2 peptides per protein group.

Presence of SNX9 in RPE-1 cell filopodia were scored by drawing a linescan region of interest over a filopodium and in an adjacent area with no filopodium in FIJI. The standard deviation in maximal intensity value of background was calculated and the numbers of filopodia with maximal intensity values more than one standard deviation above background were counted.

Value and number of N, statistical tests used, and what error bars represent can be found in the figure legends. Statistical significance of results was defined as P<0.05 (*), P<0.01 (**), P< 0.001 (***), P<0.0001 (***). Graphs were generated and statistical analysis performed in either Prism (Graphpad), R or Python using Matplotlib and Seaborn. FLS data analysis was performed with Pandas and Scipy in a Jupyter Lab environment.

## DATA AND CODE AVAILABILITY

The code for FLSAce will be available on Github.

## KEY RESOURCES TABLE

### Supplementary Figure Legends

**Sup Fig. 1.**
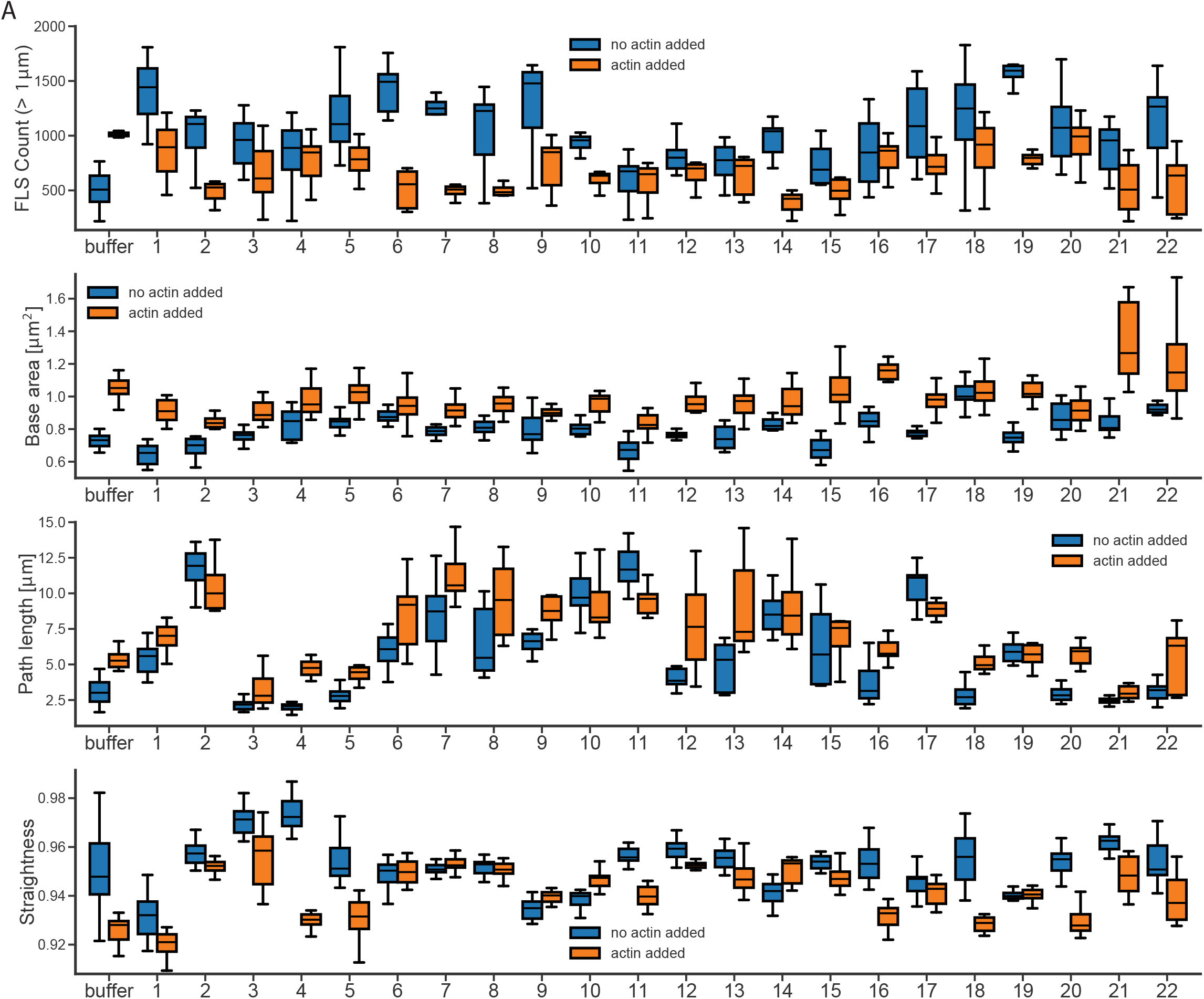
Quantification of FLS phenotypes on antibody addition. Tukey’s boxplots showing the output from FLSAce analysis for FLS assays where each different scFV is preincubated in the reaction mix (Figure 1 C-D), specifically illustrating the effects on number, base area (FLS thickness), length (path length) and straightness of each scFV. Data are the median, quartiles and range. Data are from 3 repeats for each FcFv containing no additional actin (blue bars) and 2 further repeats in which of 1 μM of supplemental unlabelled actin was added (orange bars). The addition of unlabelled actin makes FLS longer, thus increasing the length over which morphologies, such as curliness, can be seen. Additional actin also means that the phenotype of antibodies that lead to longer FLS (perhaps by inhibiting depolymerization processes) are less limited by the concentration of actin and conversely, endogenous concentrations of actin (i.e. the no additional actin condition) can make phenotypes of shorter or fewer FLS more obvious.

**Sup Fig. 2.**
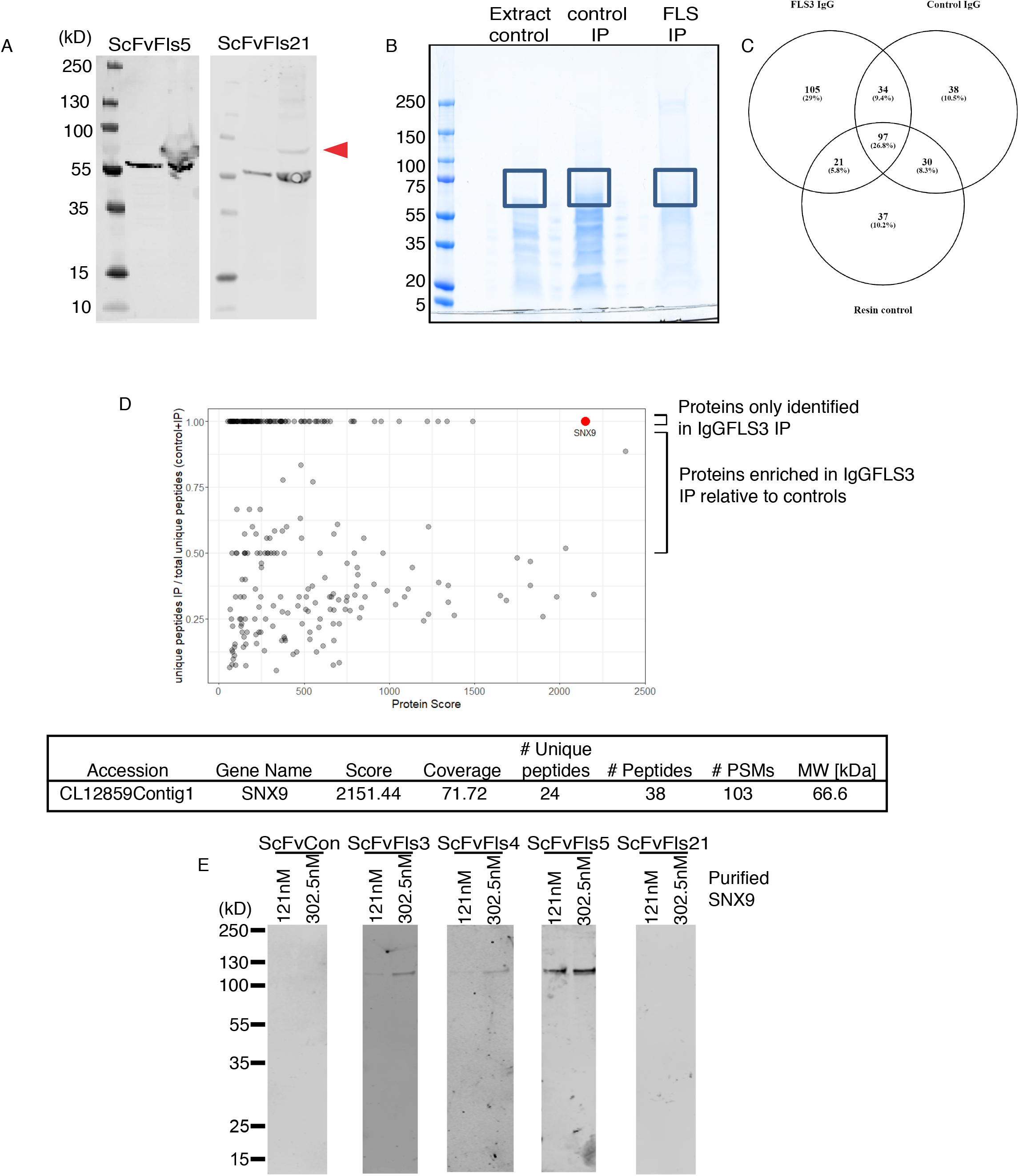
SNX9 is the antigen to scFvs 3, 4, 5, and 21. (A) Western blot of two concentrations of *Xenopus* egg extracts (for each blot left lane = 25 μg, right = 100 μg extracts) using the indicated scFvs. ScFvFls5 and 21 both detect a specific ~70 kDa band. (B) Extracts, control and IgGFLS3 immunoprecipitation (C) Venn diagram of mass spectrometry identification from the gel sectors indicated in (B). (D) SNX9 gives the highest protein score and number of unique peptides (E) Western blot of two concentrations of purified SNAP-SNX9 (for each blot left lane = 121 nM, right = 302.5nMSNX9) using the indicated scFvs, showing that scFvs 3, 4 and 5 all recognise purified SNX9, while scFv21 does not.

**Sup Fig. 3.**
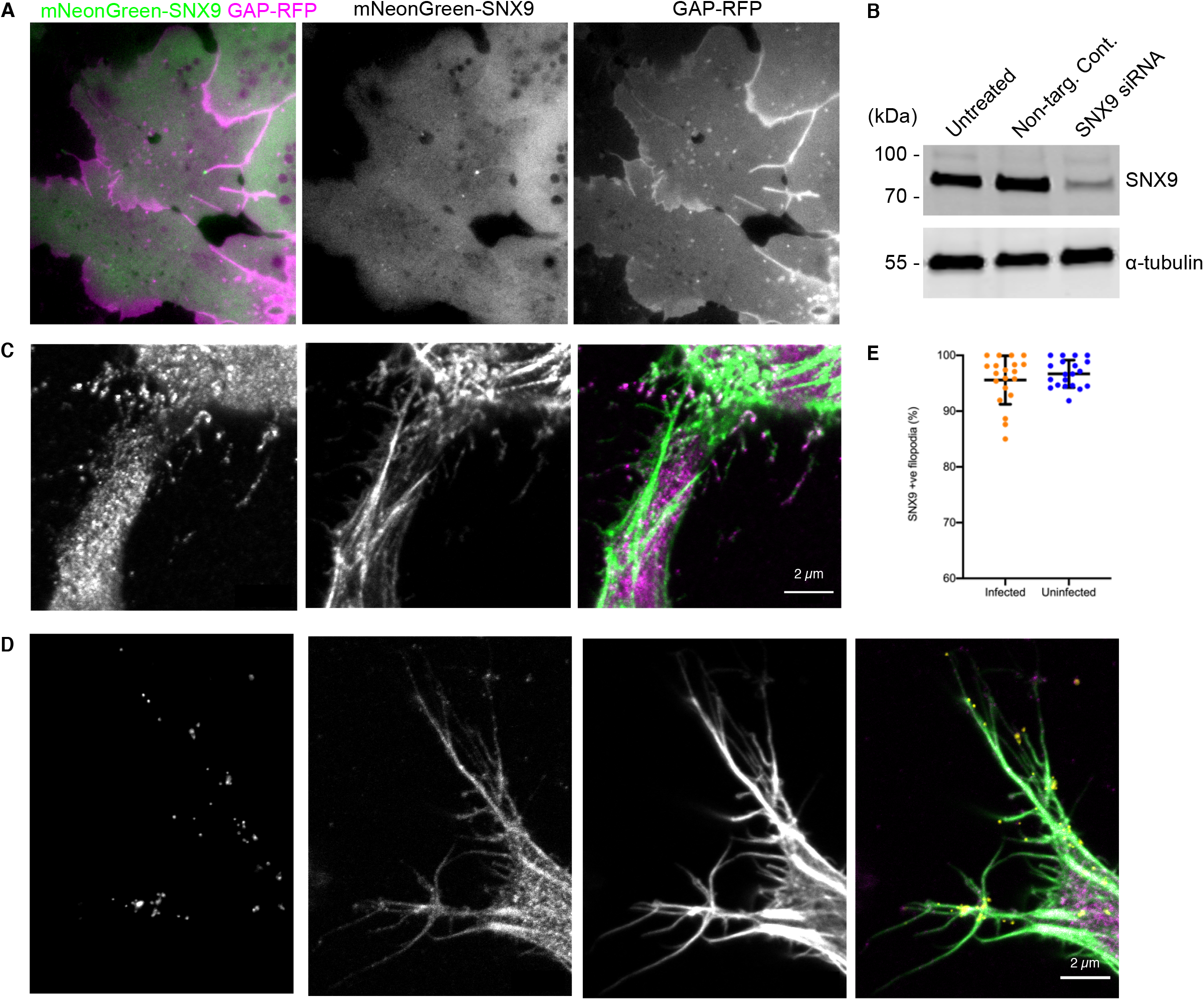
SNX9 in cells. (A) Single frame from a TIRF microscopy timelapse of *Xenopus* chordamesoderm cells expressing mNeonGreen-SNX9 (green) and GAP-RFP membrane marker (magenta) showing SNX9 at the tip of filopodia. Scale bar = 10 μm.(B) Western blots against SNX9 or the loading control DM1α (tubulin) of untreated RPE-1 cells and cells treated with either non-targeting control or anti SNX9 siRNA showing knockdown with the SNX9 siRNA sequence.

**Sup Table 1.** Statistics for effect of immunoblock and addition of SNX9 on FLS. Analysis uses a two-sample Kolmogorov-Smirnov test with Bonferroni correction for multiple comparisons.

|  | immunoblock vs control |  | rescue vs control |  | rescue vs immunoblock |  |
| --- | --- | --- | --- | --- | --- | --- |
|  | p-value | significance | p-value | significance | p-value | significance |
| scFvFLS3 | 0.000170119 | *** | 0.434495631 | n.s. | 0.00026647 | ** |
| scFvFLS4 | 3.28555E-05 | *** | 0.615688038 | n.s. | 1.39799E-05 | *** |
| scFvFLS5 | 7.26221E-06 | *** | 0.000428209 | ** | 3.25085E-05 | *** |
| scFvFLS21 | 5.36972E-05 | *** | 1.14433E-05 | *** | 0.000470845 | ** |

Sup Table 1
|  | immunoblock vs control |  | rescue vs control |  | rescue vs immunoblock |  |
| --- | --- | --- | --- | --- | --- | --- |
|  | p-value | significance | p-value | significance | p-value | significance |
| scFvFLS3 | 0.001216211 | ** | 0.916400522 | n.s. | 0.00026647 | ** |
| scFvFLS4 | 0.00026647 | ** | 0.64837074 | n.s. | 1.39799E-05 | *** |
| scFvFLS5 | 5.36972E-05 | *** | 0.735840017 | n.s. | 3.89787E-05 | *** |
| scFvFLS21 | 0.000333641 | ** | 0.000739872 | ** | 0.000333641 | ** |

**Sup Movie 1 SNX9 is active at filopodial tips in*Xenopus* dorsal marginal zone explants.** TIRF microscopy timelapse of *Xenopus* chordamesoderm cells expressing mNeonGreen-SNX9 (green) and GAP-RFP (magenta) showing active filopodia with SNX9 at their tips. Frames were captured every 2 s and are replayed at 15 fps. Scale bar = 10 μm.

---

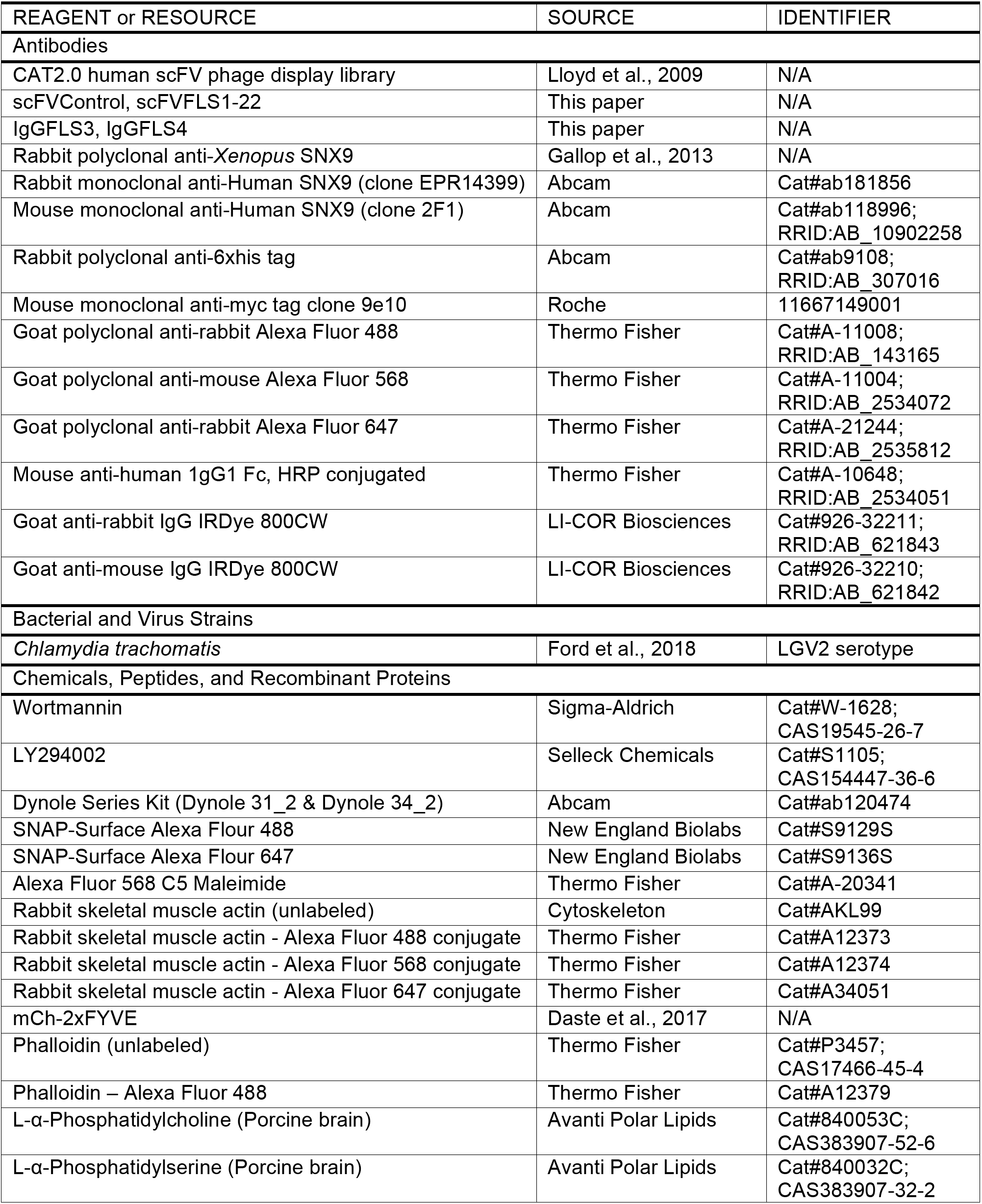
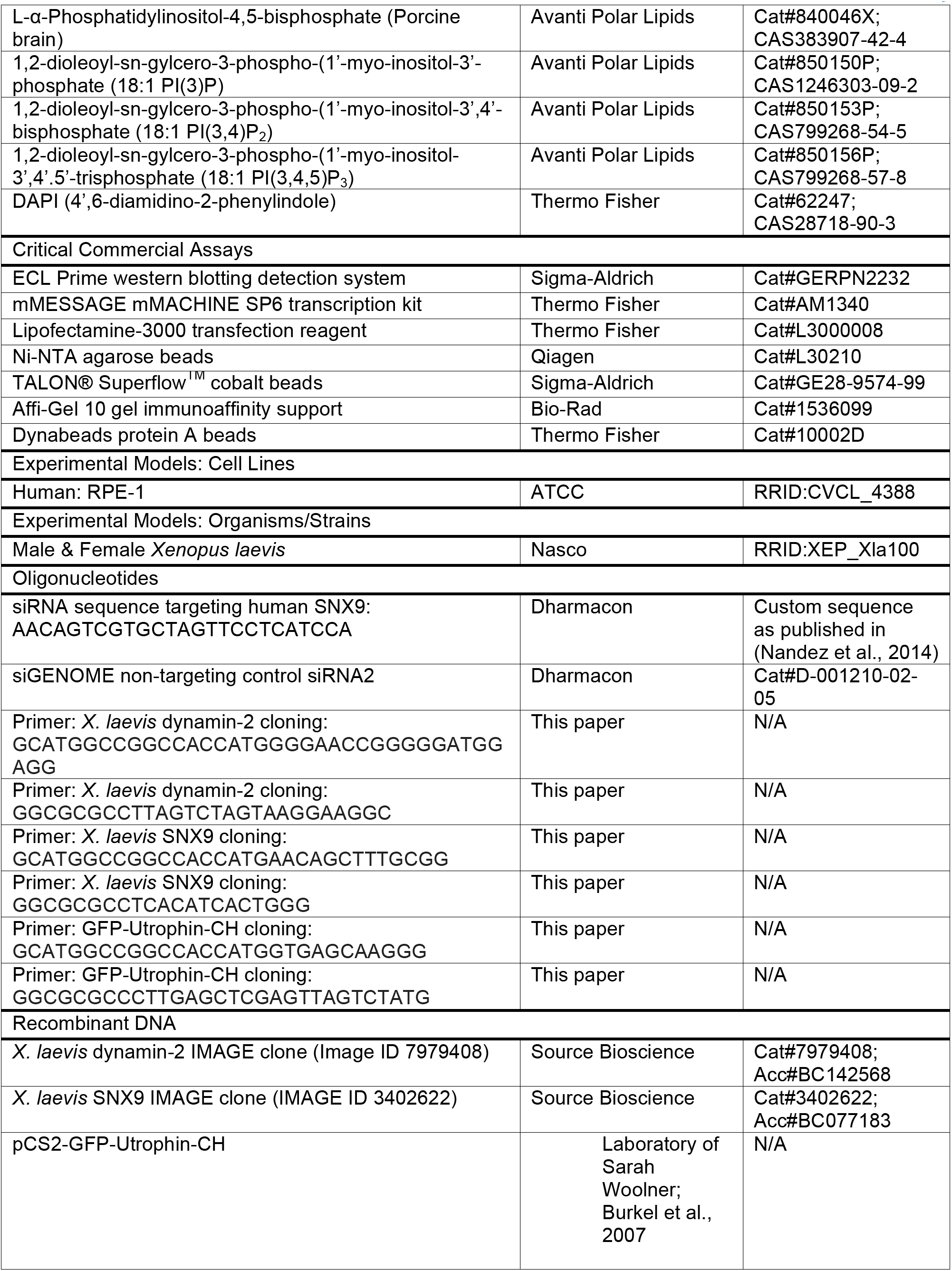
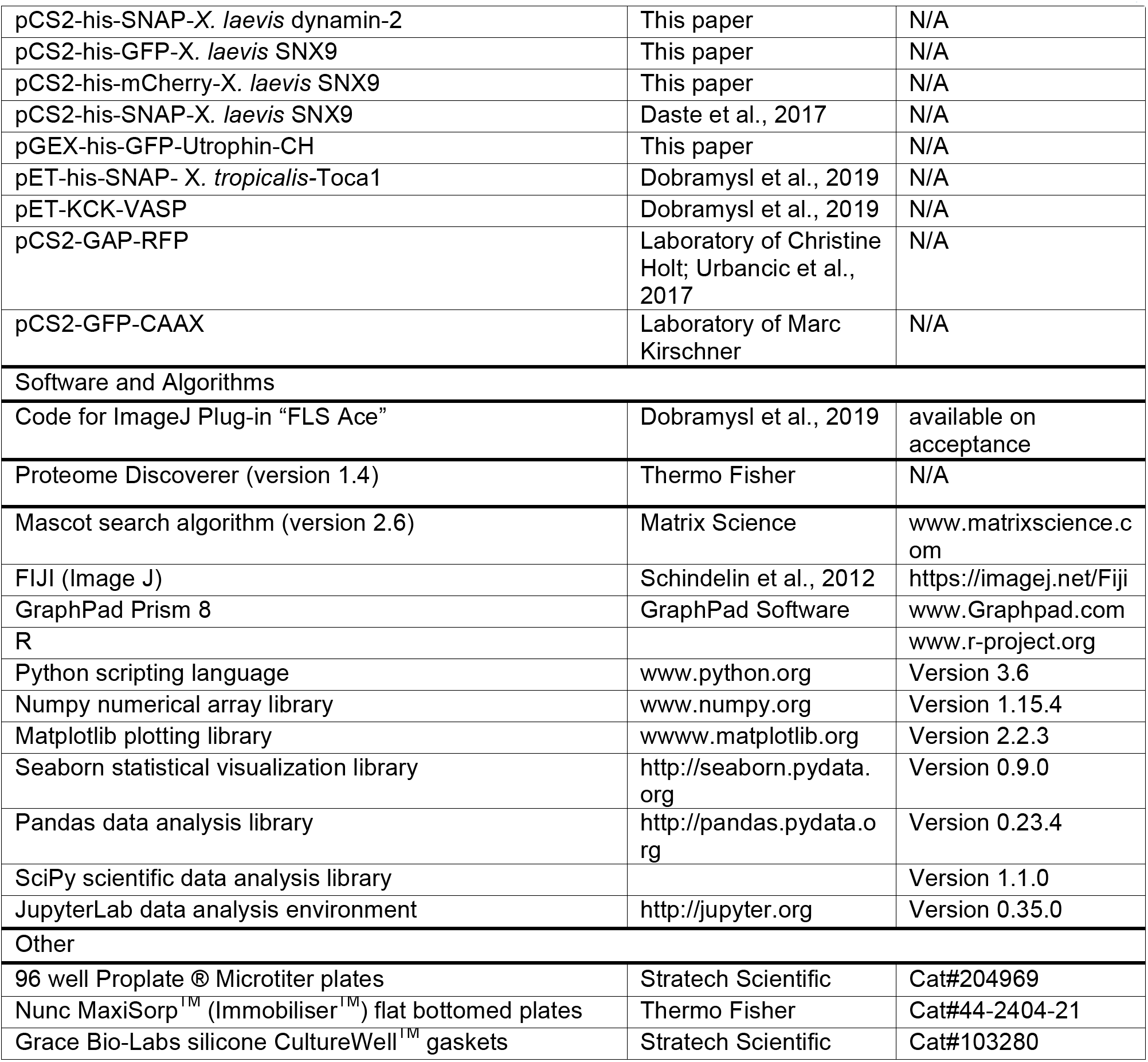
KEY RESOURCES TABLE.

